# The MYB33, MYB65, and MYB101 transcription factors affect Arabidopsis and potato responses to drought by regulating the ABA signaling pathway

**DOI:** 10.1101/2021.04.07.438025

**Authors:** Anna Wyrzykowska, Dawid Bielewicz, Patrycja Plewka, Dorota Sołtys-Kalina, Iwona Wasilewicz-Flis, Waldemar Marczewski, Artur Jarmolowski, Zofia Szweykowska-Kulinska

## Abstract

**Aims:** Drought is a climate threat limiting crop production. Potato is one of the four most important food crops worldwide and is sensitive to water shortage. The CBP80 gene was shown to affect plant response to drought by regulating the level of microRNA159, and, consequently, the levels of the MYB33 and MYB101 transcription factors. (TF) Our studies aimed to show whether indeed the level of MYB33, MYB65, and MYB101 TFs affects plant response to water shortage.

**Methods:** Arabidopsis transgenic plants exhibiting downregulation and Arabidopsis and potato transgenic plants exhibiting overexpression of selected MYB TFs were obtained. Plants response to drought was mainly measured using relative water content (RWC) and stomata closure upon exogenous ABA.

**Results:** Three MYB TFs studied are involved in plant response to drought. When downregulated in Arabidopsis, the MYB33, MYB65 and MYB101 genes cause stomatal hyposensitivity to ABA, leading to reduced tolerance to drought. Transgenic Arabidopsis and potato plants overexpressing a mutated version of these genes with changed miR159 recognition site, show hypersensitivity to ABA and relatively high tolerance to drought conditions.

**Conclusions:** The MYB33, MYB65, and MYB101 genes are good be potential targets for innovative breeding to obtain crops with relatively high tolerance to drought.

## Introduction

Drought is one of the most visible effects of climate change currently occurring worldwide. These effects are also some of the most drastic factors affecting agriculture and the economy. Via classic and molecular approaches, many genomic loci and genes potentially associated with the response to water shortage have been identified in model and crop species (Chaves et al. 2003; Duque et al. 2013; Joshi et al. 2016; Kulkarni et al. 2017). Potato (*Solanum tuberosum* L.) is considered a drought-sensitive crop species, although cultivar-dependent differences in tolerance have been described (Soltys-Kalina et al. 2016). Tolerance to drought is a very complex polygenic trait in potato (Obidiegwu et al. 2015). Several genes as well as quantitative trait loci (QTL) for drought tolerance have been identified on all 12 potato chromosomes (Anithakumari et al. 2011; Anithakumari et al. 2012; Khan et al. 2015; Pieczynski et al. 2018).

Abscisic acid (ABA) is a phytohormone that is produced in response to abiotic stress, including drought stress (Lee and Luan 2012). ABA acts at the cellular level by inducing stomatal closure and inducing/repressing stress-related genes. DREB2A/2B, AREB1, RD22BP1 and MYC/MYB represent the main group of transcription factors that regulate ABA-responsive gene expression. These proteins bind to their corresponding cis-acting elements within promoter regions, such as DRE/CRTs, ABREs and MYCRS/MYBRS (Fujita et al. 2011; Nakashima and Yamaguchi-Shinozaki 2013; Nishimura et al. 2009; Roychoudhury et al. 2013; Wang et al. 2011).

Genes involved in RNA metabolism have been found to affect the ABA signaling pathway (Hugouvieux et al. 2001; Hugouvieux et al. 2002; Laloum et al. 2018; Pieczynski et al. 2013; Reyes and Chua 2007). For example, it was found that alternative splicing, affected by SR proteins, can produce three alternative protein isoforms of the Zinc-Induced-Facilitator-like 1 (ZIFL1) transporter required for polar auxin transport in *Arabidopsis thaliana(Eckardt 2013)*. A shortened form of the transporter, ZIFL1.3, is located exclusively in the plasma membrane of leaf stomatal guard cells and is involved in drought tolerance via stomatal closure, most likely by modulating potassium and proton fluxes in plant cells.

Another group of proteins affecting the plant response to abiotic stresses is the glycine-rich RNA-binding proteins (GRPs), of which hnRNP-like proteins are members. However, the functions of these proteins remain largely unknown. In Arabidopsis, GRP7 exhibits a negative effect on plant development in response to drought stress during seed germination, seedling growth and stomatal movement (Kim et al. 2008). GRP7, as well as GRP8, binds to mRNA, which affects splicing, and affects microRNA biogenesis (Koster et al. 2014; Streitner et al. 2008).

The Arabidopsis splicing factor protein STABILIZED1, which is a homolog of the human U5 snRNP-associated protein, is involved in pre-mRNA splicing and turnover of unstable transcripts. It also affects microRNA biogenesis via pri-miRNA splicing. This protein is a regulator of plant responses to drought as well as to other abiotic stresses (Ben Chaabane et al. 2013; Kim et al. 2017; Lee et al. 2006; Shin et al. 2011).

The CAP-BINDING PROTEIN 20 (CBP20) and CAP-BINDING PROTEIN 80/ABSCISIC ACID HYPERSENSITIVE1 (CBP80/ABH1) genes were found to affect plant sensitivity to ABA and the ABA signaling pathway (Hugouvieux et al. 2001; Hugouvieux et al. 2002; Pieczynski et al. 2013). CBP20 and CBP80 form a nuclear cap-binding complex (CBC) that binds to the cap structure and affects pre-mRNA splicing, mRNA stability and microRNA biogenesis (Flaherty et al. 1997; Izaurralde et al. 1994; Kmieciak and Jarmolowski 2002; Lewis et al. 1996; Pieczynski et al. 2013; Raczynska et al. 2010; Szarzynska et al. 2009). Arabidopsis *cbp20* and *cbp80/abh1* mutants were found to be hypersensitive to ABA during germination (Hugouvieux et al. 2001; Reyes and Chua 2007).

Moreover, adult *cbp20* and *cbp80/abh1* mutants exhibit drought tolerance (Pieczynski et al. 2013). The responses concerning *CBP80/ABH1* and *CBP20* gene activity and drought tolerance are evolutionarily conserved. In crop species, inactivation of the *CBP80/ABH1* gene in potato and the *CBP20* gene in barley results in drought-tolerant phenotypes (Daszkowska-Golec et al. 2013; Pieczynski et al. 2013). A model explaining the role of the plant *CBP80/ABH1* gene in drought tolerance has been proposed. In this model, the lack of CBP80/ABH1 protein downregulated the level of miR159, which targets the mRNA of the transcription factors MYB33 and MYB101. MYB transcription factors have already been reported to function in ABA signaling and plant responses to drought (Abe et al. 1997; Casaretto et al. 2016; Jung et al. 2008; Reyes and Chua 2007; Shin et al. 2011). As a result of miR159 downregulation in *cbp80/abh1* mutant plants, increased levels of *MYB33* and *MYB101* occurred, which probably resulted in increases in MYB33 and MYB101 transcription factors. This effect was proposed as being crucial for plant tolerance to drought (Pieczynski et al. 2013).

This paper aimed to indicate the above-described model showing that the level of selected MYB transcription factors affects plant tolerance to drought. We show that downregulation of *MYB33, MYB65* and *MYB101* gene expression renders Arabidopsis plants more sensitive to drought. We added *MYB65* to our project as it was not studied as extensively as *MYB33* and *MYB101*, but shares high similarity to *MYB33*. Consequently, overexpression of these genes in Arabidopsis and potato plants resulted in increased drought tolerance. Plants with downregulated *MYB33, MYB65* and *MYB101* expression were hyposensitive to ABA and closed their stomata in response to higher ABA concentrations compared with wild-type plants. In contrast, plants overexpressing *MYB33, MYB65* or *MYB101* are hypersensitive to ABA and close their stomata in response to lower ABA concentrations than that found in wild-type plants. Thus, the lack of CBP80/ABH1 impairs miR159 biogenesis, resulting in an increase in the studied MYB transcription factors, which results in improvements to plant responses to drought.

## Material and methods

### Plant material and growth conditions

*Arabidopsis thaliana* ecotype Columbia-0 wild-type plants were used as controls and for transformation with overexpression genetic constructs. Wild-type plants; insertion mutants from the SALK collection (accession numbers: SALK_053624, SALK_042186, SALK_058312, and SALK_015891, which have a downregulated *MYB33* gene; SALK_063552, SALK_112364, and SALK_081162, which have a downregulated *MYB65* gene; and SALK_061355, which has a downregulated *MYB101* gene); and *MYB33, MYB65*, and *MYB101* overexpression (OE) transgenic Arabidopsis plants were grown in soil (Jiffy-7 42 mm; Jiffy Products International AS, Stange, Norway) in growth chambers (Sanyo/Panasonic, Japan) that had a 16-h day length (150–200 µE/m^2^s), a constant temperature of 22 °C and 70% humidity. All experiments, except those involving the selection of transformants, were conducted on 4-week-old plants and with three replicates (plants were sown at three separate times) within each from three to five biological replicates. Selection of transformants was conducted by sterilization of Arabidopsis seeds in 10% sodium hypochlorite in 70% EtOH solution, then they were grown on ½-strength MS media with 0.8% agar square plates, to which hygromycin B was added as a selective marker. Seedlings that survived were then transferred to soil, and the seeds were collected for further use. Plants of the potato cultivar Désirée were grown and propagated *in vitro* as described by Strzelczyk-Żyta (2017) in glass tubes (ø 25 mm, 150-200 mm height) filled to ∼1/5 of their height with ½ MS media with 0,1% agar; the tubes were capped with a cellulose cork and then wrapped with 1,5-2 cm wide Parafilm® M, after which they were incubated under constant light (150–200 µE/m^2^s) and a constant temperature of 22 °C. The plants were transferred from *in vitro* culture to cuvettes (35×50×12 cm) filled with soil. To mitigate stress derived from the transplantation of plants from *in vitro* culture to greenhouse conditions, the plants were kept under plastic caps for approximately 2 weeks. Afterward, the caps were removed, and the plants were grown for another 4 weeks. The plants were then transplanted to 15 cm diameter pots filled with soil and grown in a greenhouse until they were 30 – 35 cm tall. For each line, six plants of equal size were transferred to a growth chamber (16 h, 23 °C days; 8 h, 15 °C nights; light intensity above the canopy of 120 µE/m^2^s).

### Nucleic acid isolation, cDNA synthesis and PCR

Total RNA from three-week-old or four-week-old plant leaves was isolated from Arabidopsis plants and from *in vitro* potato cultures using a Direct-zol RNA MiniPrep Kit (Zymo Research). A TRIzol™ reagent (Invitrogen)-based protocol was used for the reverse transcription reaction performed with Superscript™ III Reverse Transcriptase (Invitrogen), and oligo-dT was used a primer (Szarzynska et al. 2009) in accordance with the protocol provided by the manufacturer. The RNA was then cleaned with Turbo™ DNase (Invitrogen) according to the provided protocol. Bands containing PCR products were cut from gel, and/or PCR products after the reaction were extracted from the gel and cleaned with a GeneJET Gel Extraction and DNA Cleanup Kit (ThermoFisher Scientific) according to the protocol provided by the manufacturer. The products were subsequently cloned into a pGEM®-T Easy vector (Promega). Real-time PCR was then performed with Power SYBR™ Green PCR Master Mix (Applied Biosystems) (5µl of MasterMix, 0,5 µl of each 1µM primer and 4µl of cDNA) using a 7900HT Fast Real-Time PCR System or a QuantStudio™ 7 Flex Real-Time PCR System (Applied Biosystems). The mRNA expression levels were calculated with the 2^-ΔΔCt^ method. The Mann-Whitney U test was used for statistical analyses. The following *p* values were set as statistically significant: *p*<0.05; *p*<0.01; *p*<0.001.

Genomic DNA was isolated from Arabidopsis plants for genotyping with a “fast” protocol by grinding small leaves in a 1,5 ml Eppendorf tube via a small plastic pestle and extraction buffer containing 10% SDS, EDTA, NaCl and Tris–HCl, after which the DNA was precipitated with isopropanol. Plasmid DNA was isolated with a GenElute™ Plasmid Miniprep Kit (Sigma Aldrich) according to the provided protocol.

### Genetic constructs

Constructs for overexpression were prepared with a pMDC32 Gateway™ binary plasmid containing the CaMV 35S promoter, the hygromycin B phosphotransferase gene as a selective gene, and kanamycin and rifampicin resistance genes (Curtis and Grossniklaus 2003). The cDNA sequences of the genes of interest were inserted into pENTR™/D-TOPO™ (Invitrogen) plasmids and transferred to pMDC32 using the Gateway™ LR Clonase™ II (Invitrogen) technique. The nucleotide sequences of *A. thaliana MYB33* and *MYB65* with mutated miR159 recognition site genes were kindly provided by Prof. A. A. Millar (Millar and Gubler, 2005) and were amplified using the primers kMYB33 a.thATG_F/R and kMYB65 a.th F/R (Table 1). The cDNAs of *MY0B101* from Arabidopsis (AT2G32460) and both the *MYB33* (PGSC0003DMT400058426) and *MYB65* (PGSC0003DMT400015156) sequences (the accession numbers refer to the SPUD database) from *S. tuberosum* were amplified from cDNAs using primers designed via Primer Blast (https://www.ncbi.nlm.nih.gov/tools/primer-blast/; kMYB101a.thATG_F/R; kMYB65s.tATG_F/R; kMYB33s.tATG_F/R). The miR159 recognition site was altered by directed mutagenesis using a QuikChange II Site-Directed Mutagenesis Kit (Agilent Technologies); the amino acid sequence was not changed, by the Millar group, and the primers m33F/R, m65F/R and m101F/R were used (Millar and Gubler, 2005) (Table 1). To introduce cDNA sequences into the pENTR/D-TOPO™ plasmid restriction enzyme sites for NotI and AscI, the primers kMYB33a.thRE_ATG_FLAG_F/A33_ASCI, kMYB65a.thRE_ATG_FLAG_F/A65_ASCI, kMYB101a.thRE_ATG_FLAG_F/A101_ASCI, kMYB33s.tRE_ATG_FLAG_F/S33_ASCI, and kMYB65s.tRE_ATG_FLAG_F/S65_ASCI were used (Table 1). In addition, a FLAG tag was also introduced into the 5’ sites of the given MYB cDNA sequence. The *Agrobacterium tumefaciens* AGL1 strain was used for the floral dip transformation of *A. thaliana* (Clough and Bent 1998), and the LBA4404:rif^R^ pAL4404 strain was used for *S. tuberosum* transformation (Millam 2006). The potato transformation procedure was performed as described by our group before (Wyrzykowska et al. 2016).

**Table 1.**
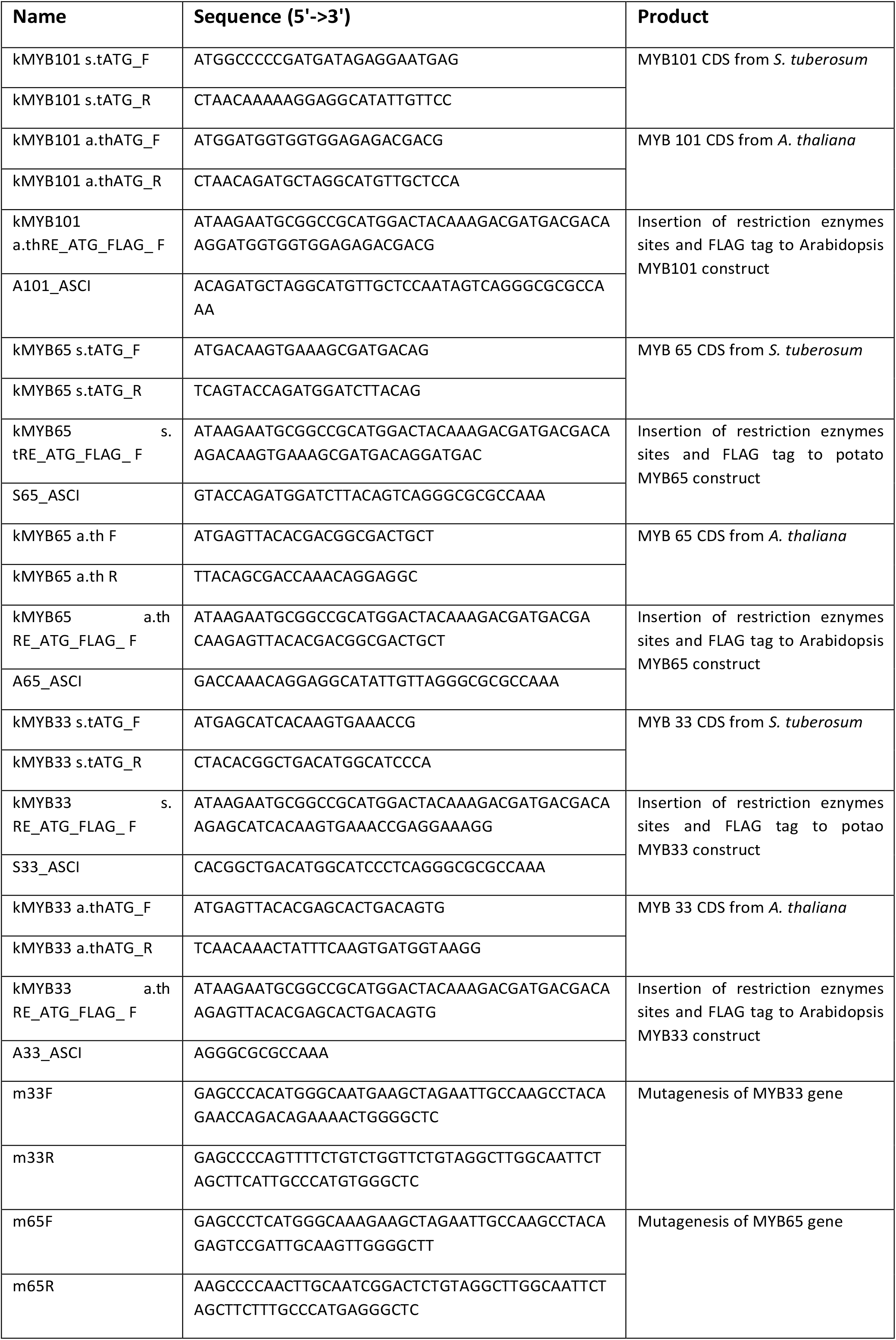
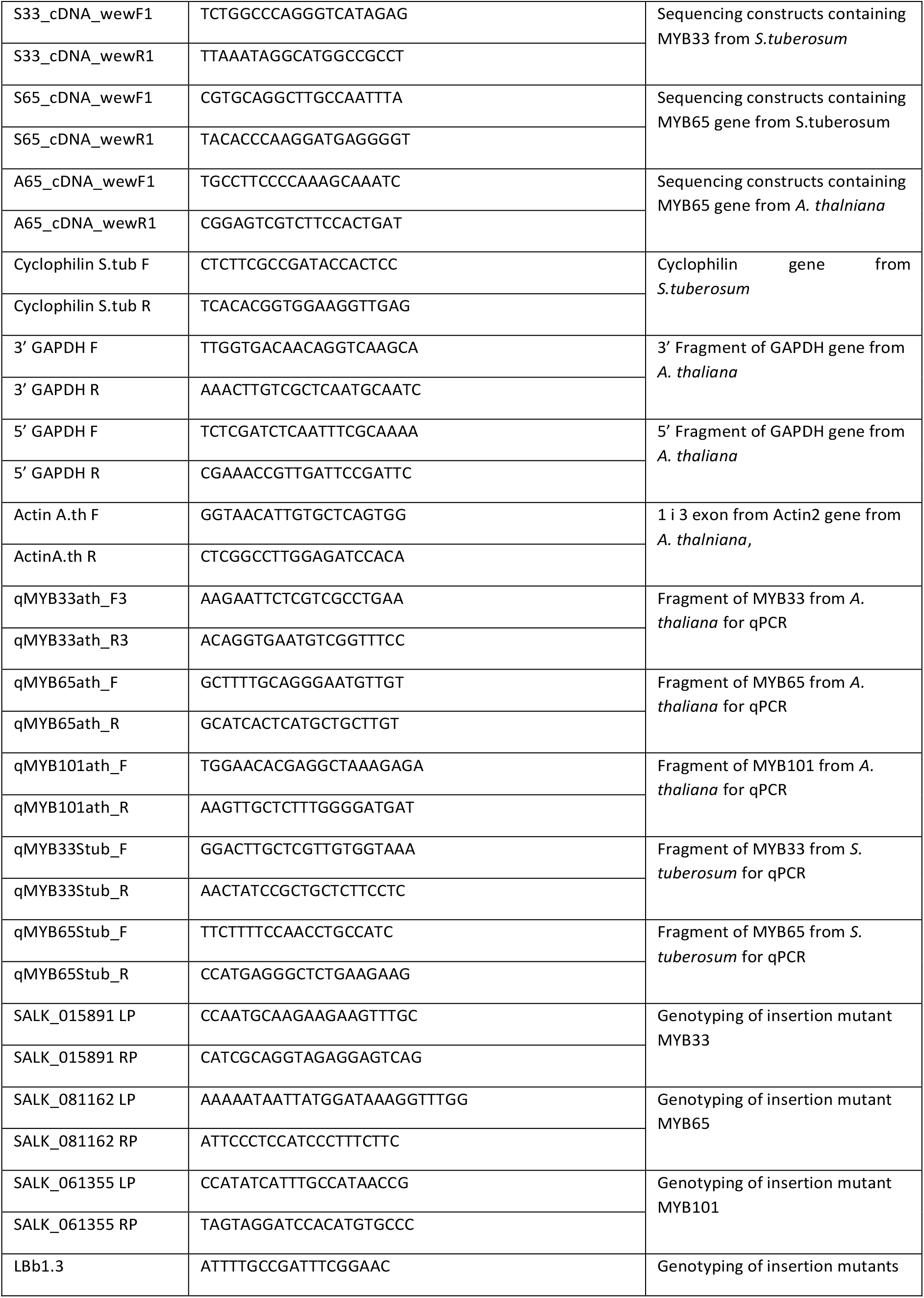
Oligonucleotides.

### Protein extraction and Western blots

Five hundred microliters of protein extraction buffer containing 200 mM Tris–HCl (pH 7.5), 250 mM NaCl, 25 mM EDTA (pH 8.0) and 0.5% SDS was added to frozen, ground plant material in an Eppendorf tube, which was subsequently vortexed at 1000-1100 rpm for 30 min at 4°C. The samples were then centrifuged at 18000 g for 30 min at 4°C. The supernatant was collected, separated into 5 tubes (100 μl per tube) and either used or stored at -20°C for a short period of time. The amount of protein in the extracts was assessed by the Bradford method using Bio-Rad Protein Assay Dye Reagent Concentrate (Bio-Rad). Protein electrophoresis in 10% or 12% PAA gels with SDS in 1x Laemmli buffer was carried out, after which the proteins were transferred to Immobilon P membranes (Merck Millipore). After blocking the membrane with 5% skimmed milk in TBS-T buffer solution for 1 h at RT or overnight at 4°C, the membrane was cut and incubated with antibodies (primary anti-actin at a 1:5000 ratio for 1.5 h at RT and anti-FLAG conjugated with horseradish peroxidase at a 1:1000 ratio for 3 h at RT). The part of membrane containing actin was then washed in TBS-T buffer 3 times for 10 min and subsequently incubated in solution containing secondary antibodies (anti-mouse at a 1:10000 ratio for 1 h at RT). After washing both parts of the membrane (as before, in TBS buffer), freshly mixed equal volumes of ECL Western Blotting Detection Reagents A and B (Amersham) were poured onto the membranes, which were subsequently incubated for 5 min and then removed. Detection of chemiluminescence was conducted via a G:Box apparatus and GeneSys (Syngene) software.

### Relative water content measurements

The relative water content (RWC) was measured as described previously (Pieczynski et al. 2013). All measurements for Arabidopsis were obtained as the average of 5 leaves that were similar in size and taken from 2 or 3 individual plants. The RWC was measured during 6 consecutive days of the drought experiment after the cessation of watering. For potato plants, RWC measurements were obtained as the average of 4 leaves that were similar in size and taken from 6 individual plants at day 0 and day 6 of drought. Statistical analysis was carried out using Mann-Whitney tests.

### Analysis of stomatal density

Experiments were carried out using fully developed rosette leaves of Arabidopsis plants. Cleared epidermal fragments peeled from Arabidopsis abaxial leaf surfaces were prepared according to the methods of Pei *et al*. and as described previously by Pieczynski et al. (Pei et al. 1997; Pieczynski et al. 2013). The adaxial side was imprinted in clear nail polish. Images of the specimens on microscope slides were taken with a Nikon Eclipse Ti light microscope and a DS-Fi1c-U2 camera, a Plan Fluor 10x DIC L N1 optical system, or a Zeiss AXIO Observer Z1 system. Counts were made using at least 30 stomata images from 3 to 5 individual plants and averaged per square millimeter of surface using NIS-Elements Advanced Research software (Nikon Instruments Europe B.V.). The number of stomata was counted on fully expanded apical leaflets from 5-week-old potato plants cultivated in the greenhouse. Three leaflets were detached from each of three plants, and adaxial and abaxial epidermal peels were removed and examined under a Nikon Eclipse E200 binocular microscope. Stomatal density was measured within an area of 1 square millimeter on three separate locations on each leaflet. Statistical analysis was carried out using the Mann-Whitney test.

### Determination of trichome density

The adaxial surfaces of *A. thaliana* leaves were examined using a binocular microscope (Leica Microsystems M60 equipped with an MC170 HD camera). Trichomes were counted using at least 30 stomata images of Arabidopsis leaves from 3 to 5 plants and averaged per square millimeter of surface using NIS-Elements Advanced Research software. The adaxial and abaxial leaflet surfaces of potato plants were evaluated for trichome density under a Nikon Eclipse E200 binocular microscope. The trichomes were counted on three 5-week-old potato plants cultivated in the greenhouse. Three leaflets were taken from each plant, and the trichome density was measured within an area of 1 square millimeter at three separate locations on each leaflet. Statistical analysis was carried out using the Mann-Whitney test.

### Stomatal movement analysis

Stomatal aperture measurements were measured as described previously (Hugouvieux et al. 2002; Pei et al. 2000; Pieczynski et al. 2013; Savvides et al. 2012). Images of the epidermal peels and measurements of stomatal aperture were analyzed in the same manner as that used for stomatal density (see above). For all ABA concentrations, at least 30 stomata images of Arabidopsis leaves from 2-3 plants and average measurements were assessed using the NIS-Elements Advance Research program (Zhang et al. 2016). Statistical analysis was carried out using the Mann-Whitney test.

### Cuticle thickness analysis

The procedure for evaluating leaf cross-sections and measuring cuticles via transmission electron microscopy was carried out according to the methods of Krzeslowska and Wozny (1996) and as previously described by Pieczynski*et al*. (2013) (Krzesłowska and Woźny 1996; Pieczynski et al. 2013). Statistical analysis was carried out using the Mann-Whitney test.

## Results

### Arabidopsis plants with a MYB33, MYB65 or MYB101 deficiency show a severe wilted phenotype upon drought and are hyposensitive to ABA

To identify Arabidopsis mutants with knocked out or knocked down expression of the investigated *MYB* genes, homozygous plants were obtained from seeds from the SALK collection (including four mutant lines with different T-DNA insertion sites for the *MYB33* gene, three lines for the *MYB65* gene and one for the *MYB101* gene-as only one is in available the collection, Fig. S1). The line with the lowest expression level of the *MYB33* gene (SALK_015891), the line with no expression of the *MYB65* gene (SALK_063552), and the line showing strong downregulation of *MYB101* expression (SALK_061355) were selected for further analyses (Fig. 1A). Four-week-old homozygous plants were subjected to drought stress by complete water cessation for 6 days, after which the RWC of the plant leaves was measured. Compared to WT plants, plants with downregulated or knocked out *MYB33, MYB 101* or *MYB65* genes exhibited a significant decrease in their tolerance to water deficit. This observation was strengthened by RWC measurements that show approximately twofold lower levels of water content in all mutant lines compared with that in the WT plants (Fig. 1B-D).

**Figure 1.**
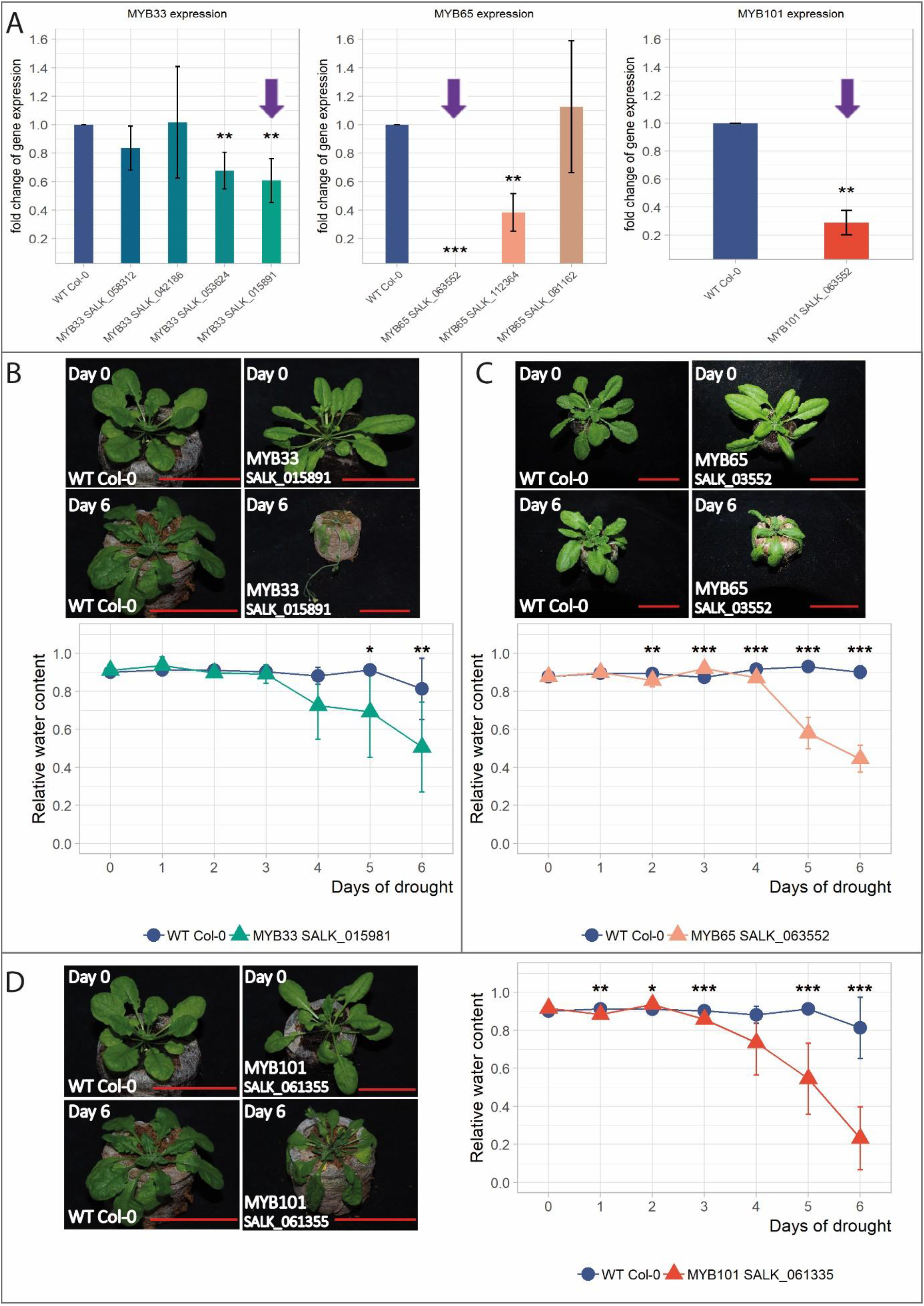
Arabidopsis plants with MYB33, MYB65 or MYB101 deficiency show a severe wilted phenotype during drought. **A)** Levels of MYB33, MYB65 and MYB101 mRNA in selected mutant plants. The left graph shows the RT-qPCR measurements of the downregulated MYB33 mRNA in four SALK lines, with line SALK_015891 exhibiting the most profound downregulation across all lines compared to that in the wild-type plants. The middle graph shows the level of MYB65 mRNA expression in three SALK lines, with the SALK_063552 line showing total knockdown of the MYB65 gene compared to that in the wild-type plants. The right graph shows MYB101 mRNA expression in the SALK_061355 mutant plant with 3.5 times lower expression compared with that in the wild-type plants. *p* value: ** *p*<0.01; *** *p*<0.001; Student t-test. The labels of the select mutant lines are shown at the bottom of each graph. The violet arrows point to the select SALK mutants used further in this study. (B, C, D) After 6 days of drought, the *myb33, myb65* and *myb101* mutant plants showed a significant wilted phenotype (upper panels) in comparison to the wild-type plants. RWC experiments (lower panels) show substantially high water loss in all three *MYB* mutant plants during the course of drought, starting on the 3^rd^ day after the cessation of watering. The images show plants on days 0 and 6 of the drought experiments involving both WT and mutants in the upper panels of B, C, and in the left panel of D. The graphs show the RWC results during the time course of drought, measured each consecutive day (blue line – wild-type plants, cyan line – *myb33* mutant plants (B), salmon line – *myb65* mutant plants (C), orangered line – *myb101* mutant plants (D)). The data are shown as the means± SDs of n=3 independent experiments; Mann-Whitney test, *p* value: * *p*<005; ** *p*<0.01; *** *p*<0.001. Scale bar – 50 mm.

These results reinforced the hypothesis that the levels of the *MYB33, MYB65* and *MYB101* genes are involved in the plant response to drought. To uncover the reasons behind the strong wilting of the mutant plants with the downregulation of the studied MYB transcription factors (TFs), we investigated selected morphological and physiological traits that often reflect plant strategies for coping with water deficiency stress, and shown in the case *of cbp80/abh1* Arabidopsis and potato mutants, which were strongly affected. These traits include stomatal (on both sides of the leaf blades) and trichome density, cuticle thickness and stomatal response to ABA. In our previous studies, a lack of *CBP80/ABH1* gene expression led to increased drought tolerance, downregulation of stomatal density on the upper leaf blades, increased numbers of stomata on the bottom leaf blades, increased numbers of trichomes on the upper leaf blades, increased cuticle density, and hypersensitivity to ABA-induced stomatal closure (Pieczynski*et al*., 2013). In the case of the *myb33, myb65*, and *myb101* Arabidopsis mutant plants, we expected either no effects or the opposite effects as those observed for the *cbp80/abh1* mutants in our previous studies. There were no clear effects on stomatal or trichome numbers or cuticle density (Fig. S2 and Fig. S3). However, after a saturating humidity treatment, the fully opened stomatal aperture was smaller in the transgenic plants than in the wild-type plants (13% in the case of the *myb33* mutant plants, 20% in the case of the *myb65* mutant plants, and 7,8% in the case of the *myb101* mutant plants). Moreover, the stomata of the mutant plants with downregulated expression of the *MYB33, MYB65* and *MYB101* genes remained open even under 10 µM ABA concentrations, while the WT plants started closing their stomata in response to concentrations as little as 0.1 µM ABA until they were completely closed in response to 10 µM ABA (Fig. 2). Thus, the *myb33, myb65*, and *myb101* Arabidopsis mutant plants exhibited hyposensitivity to ABA with respect to stomatal closure, which is in agreement with our expectations concerning the action of CBP80/ABH1 upstream of the regulatory pathway controlling ABA-dependent stomatal closure.

**Figure 2.**
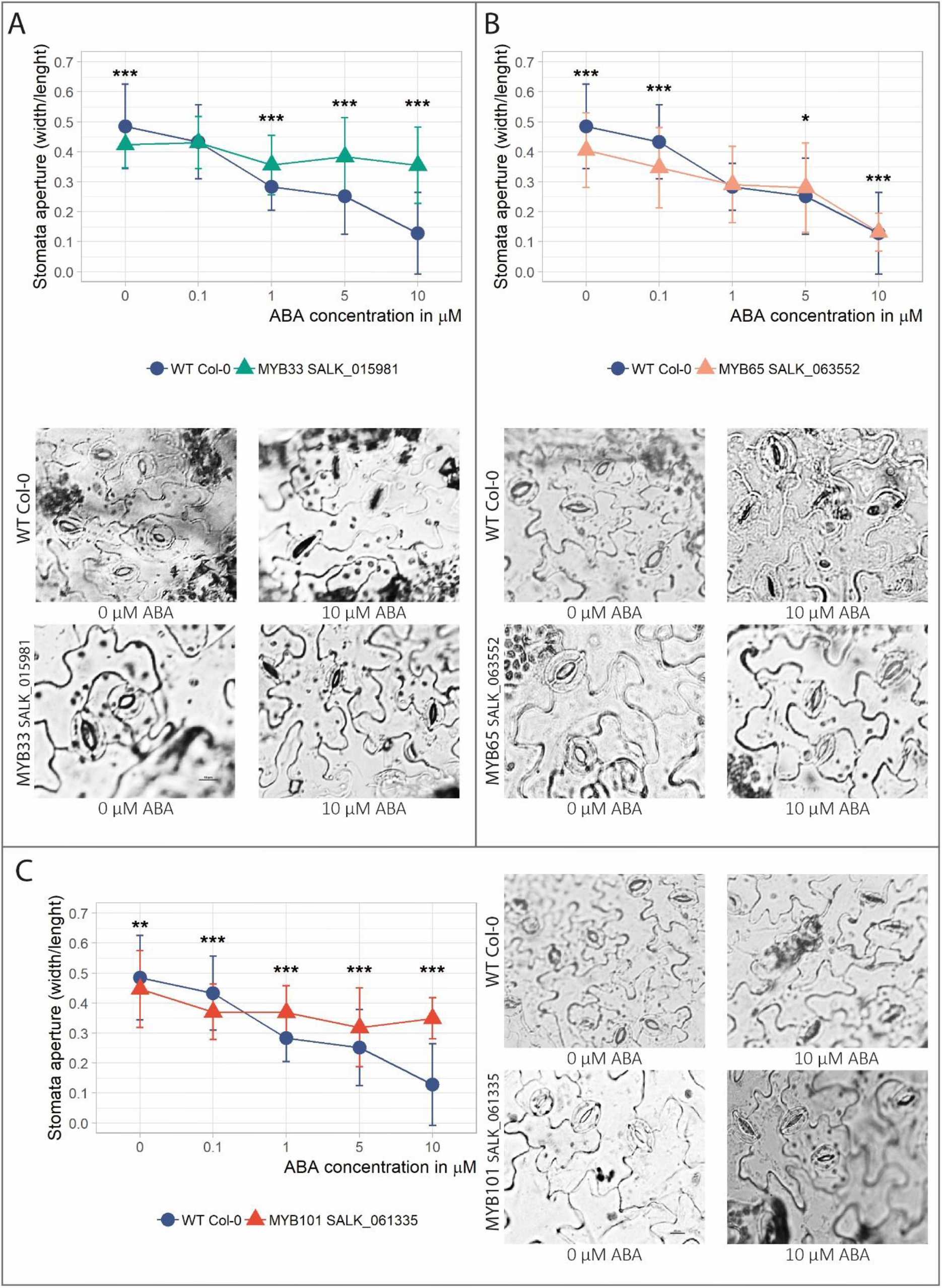
Arabidopsis mutant plants exhibiting MYB33, MYB65, or MYB101 deficiency show an ABA-hyposensitivity phenotype. **A)** Stomatal closure in ABA-hyposensitive mutant plants with MYB33 deficiency compared to wild-type plants. The wild-type and mutant plants were treated with increasing concentrations of ABA (upper panel: the graph shows the stomatal aperture). WT - blue line, *myb33* mutant – cyan line. (**B, C**) Stomatal closure in ABA-hyposensitive mutant plants with a MYB65 (B) and MYB101 (C) deficiency compared to wild-type plants. WT – blue line, *myb65* – salmon line, *myb101* – orangered line. The description of the upper and lower panels is the same as that in the case of the *myb33* mutant. (A, B, C) The lower panels show light micrographs of the stomata in the WT and *myb33, myb65*, and *myb101* mutant plants treated with 0 and 10 µM ABA. The data are shown as the means± SDs of n=3 independent experiments, with 30 stomata per data point. Scale bar – 10 µm; Mann-Whitney test, *p* value: * *p*<0.05; ** *p*<0.01; *** *p<*0.001;.

### *Arabidopsis thaliana* transgenic plants overexpressing the *MYB33, MY65*, and *MYB101* genes from Arabidopsis and the *MYB33* gene from *S. tuberosum* are tolerant to drought and hypersensitive to ABA

To determine whether Arabidopsis plants overexpressing the *AtMYB33, AtMYB65 and AtMYB101* genes exhibit drought-tolerant phenotypes, we cloned the appropriate cDNAs into a pMDC32 binary plasmid containing both the CaMV 35S promoter and the hygromycin B phosphotransferase gene as a selective marker. Additionally, a FLAG tag was added to the 5’ end of each coding sequence. The same constructs were prepared in conjunction with *MYB33* and *MYB65* cDNAs from potato (*StMYB33* and *StMYB65*). To our knowledge, there is no *MYB101* gene in the potato genome. To avoid posttranscriptional silencing of the introduced MYB transcripts by microRNA 159, the microRNA 159 recognition site of the overexpressed Arabidopsis and potato cDNAs was mutated at the nucleotide level, ensuring that the amino acid sequences were unaffected (Millar and Gubler, 2005) (Fig. S5A). The constructs were subsequently introduced into Arabidopsis wild-type plants, and homozygous plants were selected. In the case of the *AtMYB101* Arabidopsis transgenic plants, we were unable to obtain homozygous plants. We therefore analyzed the siliques of the *AtMYB101* OE transformants. The transformants showed empty spaces where seeds should develop and/or they contained underdeveloped seeds compared to WT siliques, which contained a septum fully stacked with properly developed green seeds (Fig. S4). These results clearly indicate that overly high AtMYB101 gene overexpression is embryo lethal. We decided to continue our studies on the role of AtMYB101 in plant drought tolerance using heterozygous plants, which were genotyped before each experiment. We also did not obtain Arabidopsis plants overexpressing potato *MYB65*. The expression of transgenic cassettes in select Arabidopsis transgenic lines was tested using RT-PCR and Western blotting (Fig. S5B, C). For subsequent experiments, we selected three transgenic lines with the highest expression of each introduced transgene (A33 2-1, A33 6-4, A33 6-6, A65 3-1, A65 3-2, A65 5-4, A101 2-2-2, A101 2-11-2, A101 2-11-4, S33-1, S33-5, S33 3-11).

Four-week-old Arabidopsis plants from the selected transgenic lines were subjected to drought stress by complete water cessation for 5 days, and the RWC of the leaves was measured. In each case, plants carrying the overexpression MYB construct showed significant improvement in drought tolerance compared to wild-type plants. This observation is supported by RWC measurements that show higher levels of water content in all mutant lines than in WT plants (Fig. 3 A-D). We then examined the responsiveness of stomatal closure to ABA in all mutant plants and compared it to that of the wild-type plants. After a saturating humidity treatment, the stomatal apertures were smaller in all transgenic plants than in the wild-type plants whose stomata were fully opened (∼17%, ∼30%, ∼24%, and ∼27% for *AtMYB33* OE lines, *AtMYB65* OE lines, *AtMYB101* OE lines and *StMYB33* OE lines, respectively). Furthermore, for all four genetic constructs, the plants exhibited significantly enhanced stomatal closure in response to ABA, compared to that of the wild-type Arabidopsis plants (Fig. 4 A-D). Thus, overexpression of *AtMYB33, AtMYB65, AtMYB101* and S*tMYB33* in *A. thaliana* plants resulted in hypersensitivity to ABA. These results are in full agreement with those of our previous studies on the role of *CBP80/ABH1* in the plant response to drought, emphasizing the upstream action of this protein in the ABA-dependent signaling pathway, which leads to the regulation of *MYB33, MYB65* and *MYB101* expression (Pieczynski et al. 2013).

**Figure 3.**
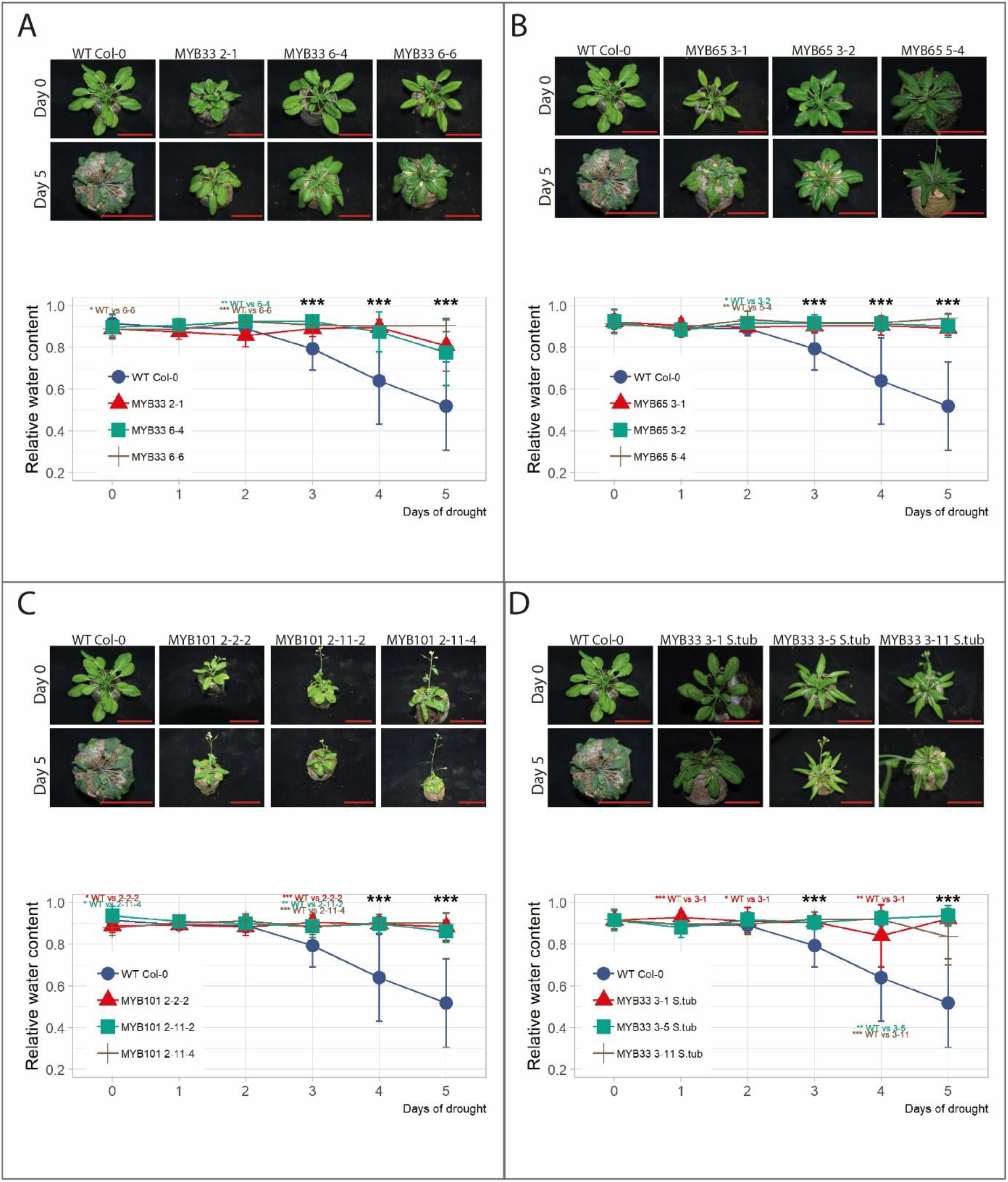
*Arabidopsis* transgenic plants overexpressing the *MYB33, MY65*, or *MYB101* gene from Arabidopsis and the *MYB33* gene from *S. tuberosum* are resistant to drought. **A)** The upper panel shows plants from three independent transgenic Arabidopsis lines overexpressing the *MYB33* gene (MYB33 OE) compared to WT plants. The plants are shown on day 0 (Day 0, upper row) and day 5 (Day 5, lower row) after water cessation. The lower panel graph shows the RWC during the time course of drought, measured each consecutive day (blue line – wild-type plants, other colored lines – plants from select independent transgenic lines overexpressing the *MYB33* gene). **(B, C)** The same experiments were performed for the *MYB65* and *MYB101* overexpression mutants. (**A, B**, and **C**) The RWC data are shown as the means± SDs of n=3 independent experiments; Mann-Whitney test, *p* value: * *p<*0.05; ** *p*<0.01; *** *p*<0.001. Scale bar – 50 mm.

**Figure 4.**
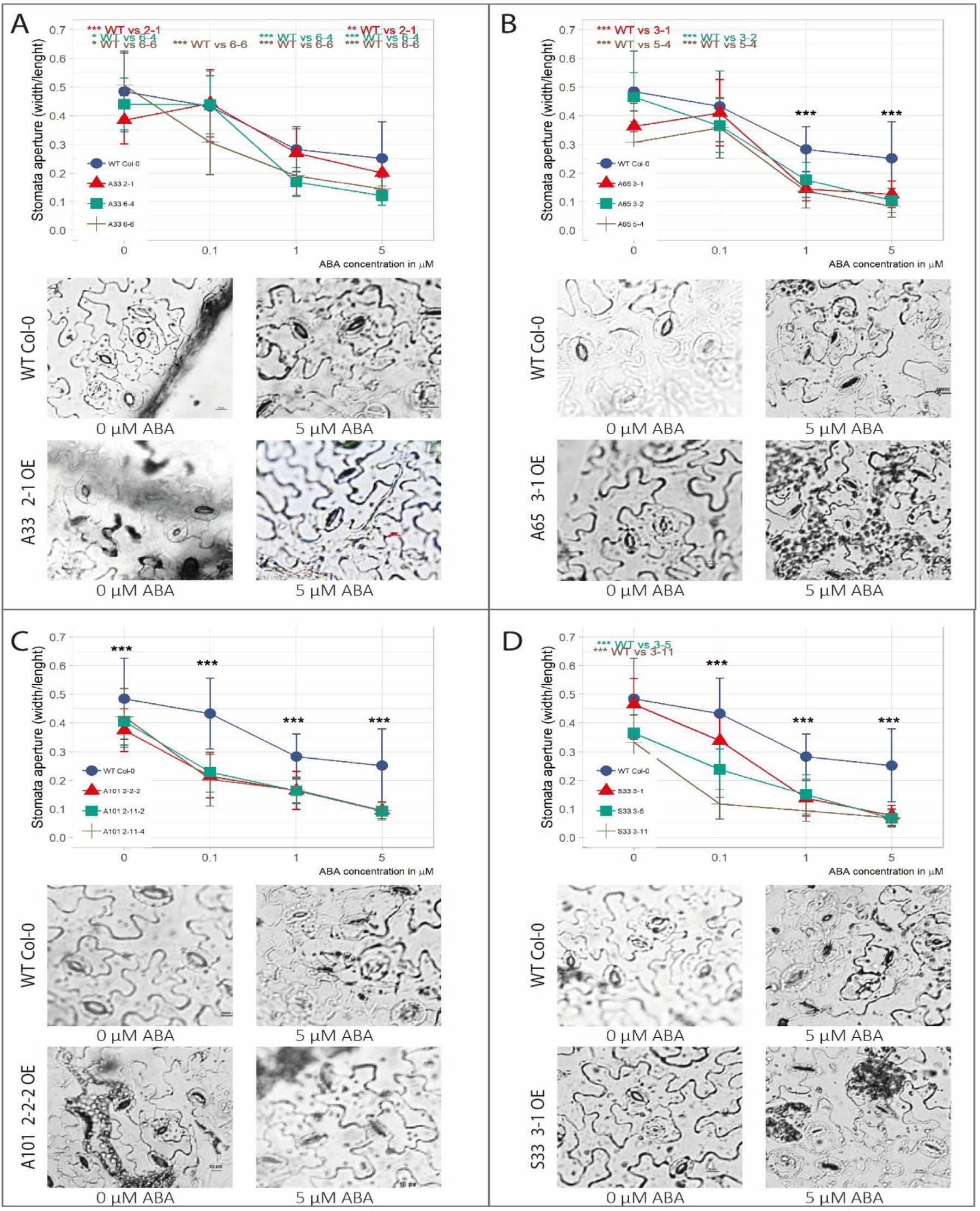
*Arabidopsis* mutants overexpressing the *MYB33, MY65*, or *MYB101* gene from Arabidopsis and the *MYB33* gene from *S. tuberosum* show an ABA-hypersensitive phenotype. **A)** Stomatal closure in ABA-hypersensitive mutant plants with *MYB33* overexpression compared to wild-type plants. The wild-type and mutant plants were treated with increasing concentrations of ABA (upper panel: the graph shows the stomatal aperture). WT - blue line, other colors – select transgenic mutant plants overexpressing *MYB33*. The lower panel shows light micrographs of the stomata in the WT plants and *MYB33* OE mutants treated with 0 and 5 µM ABA, respectively. (**B, C**) Stomatal closure in ABA-hypersensitive mutant plants with *MYB65* (B) and *MYB101* (C) overexpression compared to wild-type plants. WT – blue line, other colors – select transgenic mutant plants overexpressing *MYB65* (B) or *MYB101* (C). The description of the upper and lower panels is the same as that in the case of the *MYB33* OE mutant. (A, B, and C) The stomatal closure measurement data are shown as the means± SDs of n=3 independent experiments, with 30 stomata per data point.; Mann-Whitney test, *p* value: * *p*<0.05; ** *p*<0.01; *** *p*<0.001. Scale bar – 10 µm.

Interestingly, the numbers of stomata on the adaxial and abaxial sides of leaf blades as well as trichome density were affected in all mutant plants overexpressing MYB TFs in the same way as that in the case of the *cbp80/abh1* mutants. When the abaxial leaf surfaces were inspected, the number of stomata on the Arabidopsis mutants showed a general increase in comparison to those on the wild-type plants, although this increase was not always statistically significant. The opposite trend, a statistically significant decrease in stomata number, was observed in the case of adaxial leaf surfaces in a majority of mutant lines (Fig. S6A). The number of trichomes on the adaxial surface of leaf blades also showed a statistically significant increase in the majority of mutant lines compared to wild-type plants (Fig. S6B). In the case of the cuticle, we did not observe any changes in its density; however, it was thinner in the majority of the mutant lines than in the WT plants (Fig. S7). Thus, overexpression of the *MYB33, MYB65*, and *MYB101* genes suggests that stomatal and trichome density are at least partially under the control of the signaling pathway involving these genes and *CBP80/ABH1*, which acts upstream, while cuticle thickness is not.

### *Solanum tuberosum* plants overexpressing MYB TFs show improved tolerance to drought

A previous study on the role of CBP80/ABH1 in the plant response to drought revealed its evolutionary conservation between Arabidopsis and potato. To determine whether the role of select MYB TFs in the drought response is also conserved, we introduced the same genetic constructs containing the *MYB33, MYB65* and *MYB101* genes from *A. thaliana* and *MYB33* and *MYB65* from *S. tuberosum* into plants of potato cultivar Désirée. After Agrobacterium-mediated transformation via callus induction *in vitro*, selection of transformed tissue via cultivation on selective media, and regeneration of transgenic plants, we obtained 29 transgenic potato plants, which were transferred and cultivated in glass tubes: 12 *AtMYB33* OE plants, 6 *AtMYB65* OE plants, 4 *AtMYB101* OE plants, 6 *StMYB33* OE plants, and 1 *StMYB65* OE plant. The expression of each transgene was confirmed using RT-PCR and expression of the hygromycin B phosphotransferase gene as a control (Fig. S8 A, B). Three transgenic lines with the highest overexpression of each MYB TF studied were selected and transplanted into pots after adaptation to *ex vitro* conditions. Potato plants overexpressing *StMYB33* were smaller than wild-type potato plants. In the case of *StMYB65* OE plants, only one line was obtained. The *StMYB65* OE plants *in vitro* and those grown in pots showed a strong phenotype: they were bushy, extremely dwarfed, with shortened internodes in the stems and miniature leaves (Fig. S9). Because of these morphological defects, we decided not to include the *StMYB65* OE plants in subsequent studies.

After the transgenic potato plants grew to 30-35 cm in height, they were subjected to drought stress by complete water cessation for 6 and 11 days, and the RWC of the plant leaves was measured. In each case, the plants carrying the overexpression MYB construct showed significant improvement to drought compared to the wild-type plants. This observation was supported by the RWC measurements, which showed higher levels of water content in all the mutant lines compared with the WT lines on the 6^th^ day of drought (Fig. 5A-E). However, plants experiencing 11 days of drought showed significantly increased water contents only in the *AtMYB101* and *StMYB33* lines (Fig. S10). We also examined the responsiveness of stomatal closure to ABA in all potato OE plants and compared it to that of the wild-type plants. The experiments were conducted in the same manner as that in the case of Arabidopsis transgenic and wild-type plants. Two out of three transgenic potato lines for each MYB TF subjected to drought were used for the ABA sensitivity experiment. After a saturating humidity treatment, stomatal apertures were smaller in all the transgenic potato plants than in the wild-type plants whose stomata were fully opened (∼30%, ∼37%, ∼47%, and ∼55% for *AtMYB33*, A*tMYB65, AtMYB101* and *StMYB33*, respectively), as occurred in the case of Arabidopsis transgenic plants overexpressing the studied MYB TFs. However, for all four genetic constructs, the potato plants exhibited significantly enhanced stomatal closure in response to ABA treatment, compared to that of the wild-type potato plants (Fig. 6A-D). Our experiments clearly show the evolutionary conservation of the function in response to drought of all MYB TFs studied in plants.

**Figure 5.**
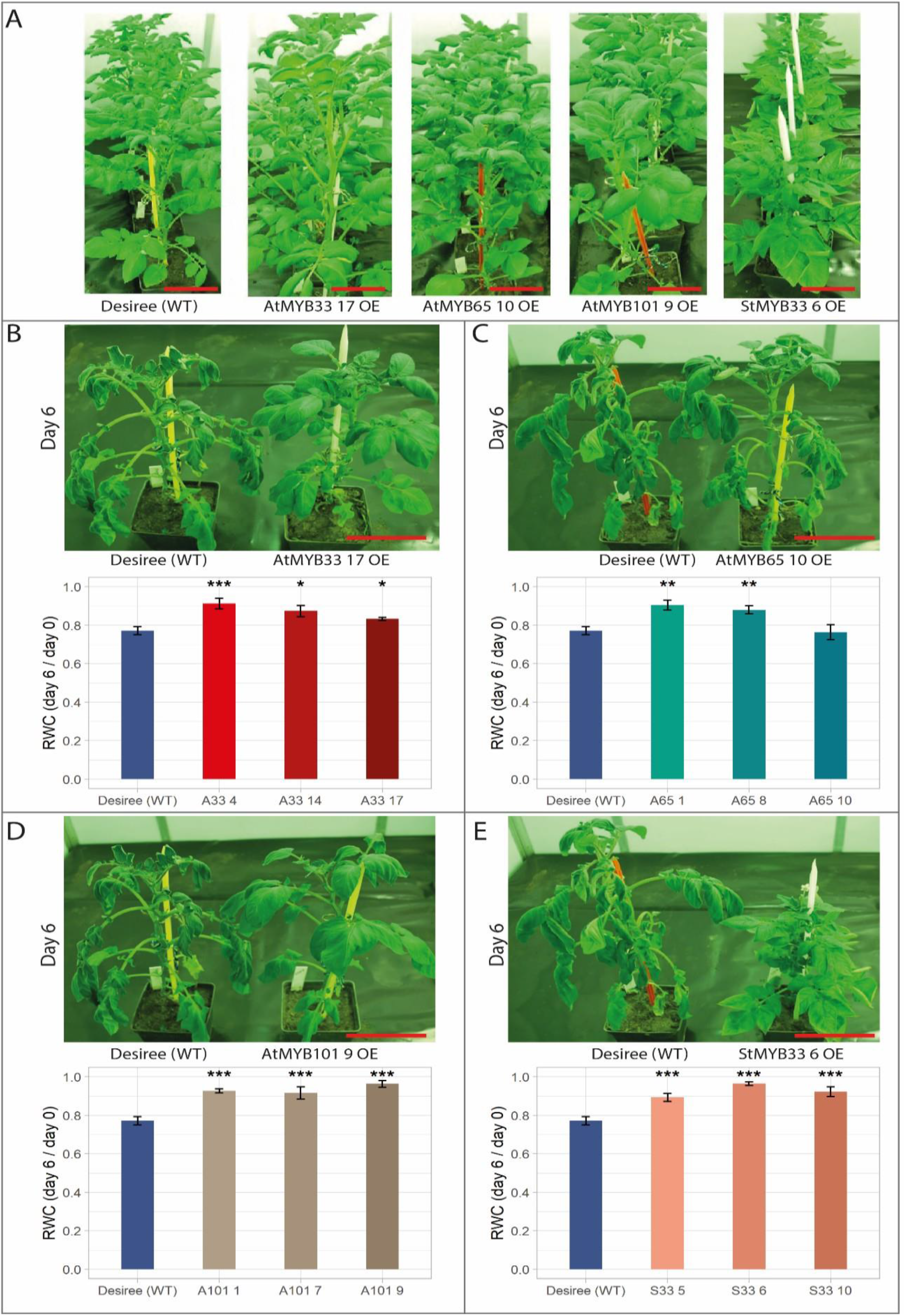
Transgenic lines of potato cultivar *S. tuberosum* cv. Désirée transgenic plants overexpressing the *MYB33, MYB65*, or *MYB101* gene from Arabidopsis and the *MYB33* gene from *S. tuberosum* are resistant to drought. (**A**) Wild-type and mutant plants overexpressing *AtMYB33, AtMYB65, AtMYB101*, and *StMYB33* on day 0 of drought. The labels shown at the bottom of each image represent the name of an individual transgenic line. (**B**) The upper panel shows transgenic potato plants overexpressing the *AtMYB33* gene (line AtMYB33-17 OE) compared with WT plants. The plants are shown on day 6 after water cessation. The lower panel shows the RWC measurements of leaves from WT and transgenic *AtMYB33* OE (A33) plants after drought stress. The RWC value of the control (Désirée – 0.77) is the value of the ratio between its water content on day 6 and on the day 0. Blue bars – WT plants, the other colored bars represent potato plants from independent transgenic lines overexpressing *AtMYB33*. (**C, D**, and **E**) show the same data obtained for potato transgenic plants overexpressing *AtMYB65* (**C**), *AtMYB101* (**D**), and *StMYB33* (**E**), respectively. The figure descriptions are the same as the description in panel (A). (B, C, D, and E) The RWC data are shown as the means± standard error of the mean, n>6 independent experiments; Mann-Whitney test, *p* value: * *p*<0.05; ** *p*<0.01; *** *p*<0.001 test. Scale bar – 15 cm.

**Figure 6.**
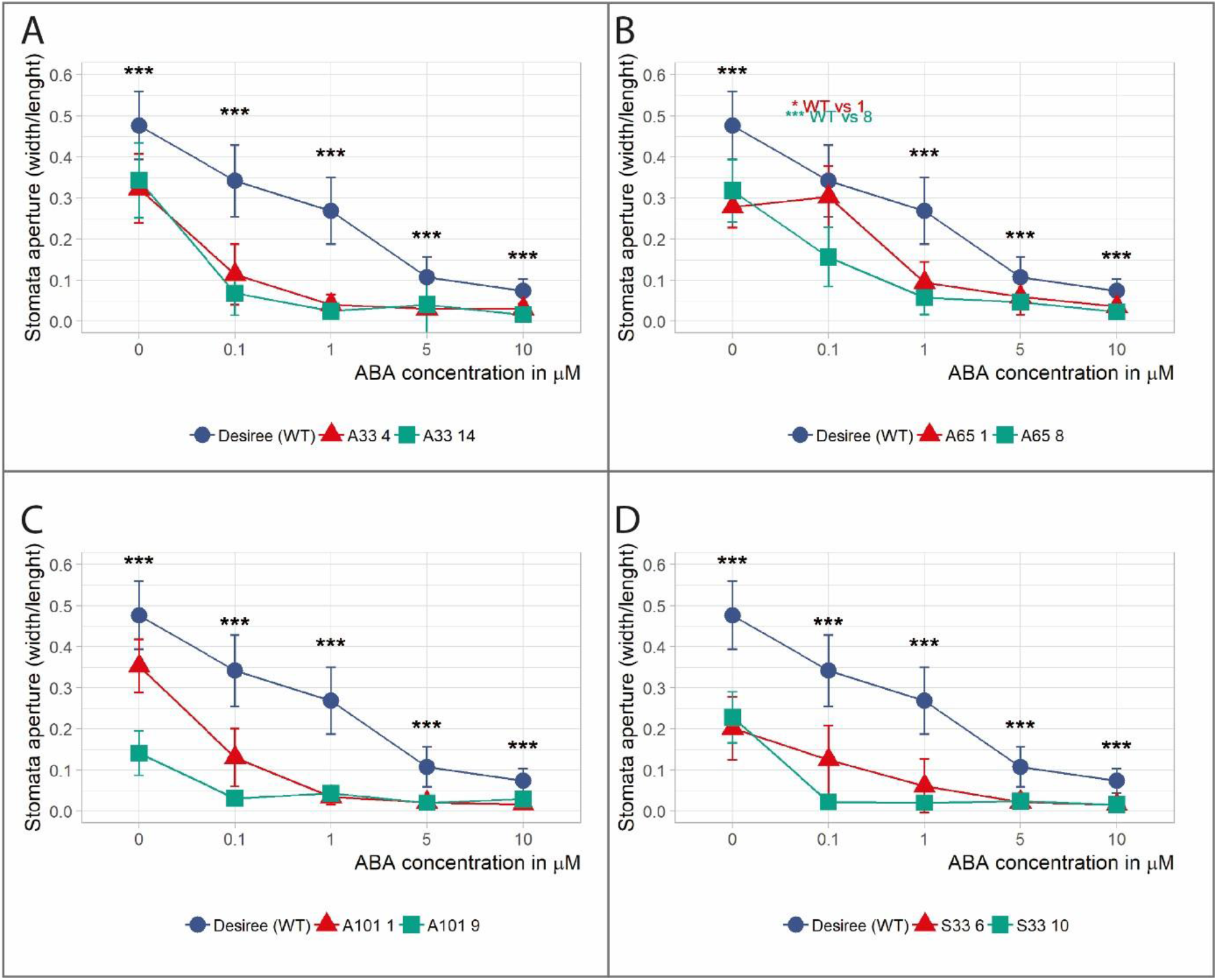
Transgenic lines of potato cultivar Désirée overexpressing the *AtMYB33, AtMYB65*, and *AtMYB101* genes and the *StMYB33* gene exhibit ABA hypersensitivity. **A)** Stomatal closure in ABA-hypersensitive transgenic potato plants with At*MYB33* overexpression compared to wild-type plants. The wild-type and mutant plants were treated with increasing concentrations of ABA. WT - blue line, other colors – select transgenic mutant lines overexpressing *AtMYB33*. **(B, C, D)** Graphs of the stomatal aperture of transgenic potato plants overexpressing *AtMYB65* (B), *AtMYB101* (C) and *StMYB33* (D). The figure descriptions are the same as the description in panel (A). The data are shown as the means± SDs of n=3 independent experiments, with 30 stomata per data point; Mann-Whitney test, *p* value: * *p*<0.05; ** *p*<0.01; *** *p*<0.001. Stomatal aperture was measured in two out of three independent transgenic lines for all overexpression constructs.

Interestingly, the overexpression of the *MYB33, MYB65* and *MYB101* genes in potato plants did not affect stomatal or trichome density in the same manner in all the transgenic lines, which is in contrast to that which occurred in the Arabidopsis plants overexpressing MYB TFs (Fig. S11).

## Discussion

The results of this work support the model of the plant drought tolerance pathway that is induced by the lack of the *CBP80/ABH1* gene presented in our previous study (Pieczynski et al. 2013). As shown previously, the lack of this gene impairs miRNA159 induction, which is known to control *MYB33, MYB65*, and *MYB101* gene expression in Arabidopsis (Alonso-Peral et al. 2010; Millar and Gubler 2005; Reyes and Chua 2007). The lack of miRNA159 induction resulted in elevated levels of *MYB33*/*MYB101* mRNAs, which purportedly increase plant tolerance to drought (Pieczynski et al. 2013). As expected, our recent results show that downregulation of *MYB33*/*65*/*101* expression resulted in decreased plant tolerance to drought, while overexpression of the same genes led to improved plant responses to water shortage. Moreover, the results of the present study show that plants with downregulated expression of MYB33/65/101 genes were hyposensitive to ABA and closed their stomata at ABA concentrations greater than wild-type plants. The opposite was observed in the case of plants overexpressing these MYB transcription factors. The role of MYB 33/101 transcription factors in ABA signaling has already been proven for seed germination in the case of Arabidopsis (Reyes and Chua 2007). However, the role of these transcription factors and MYB65 in the drought response has never been tested. Interestingly, the downregulation or upregulation of each individual MYB33/65/101 transcription factor resulted in a strong response to drought, suggesting that these proteins are not redundant in their functions related to the response to water deficit. MYB33/65/101 belong to the same R2R3 subgroup of the MYB transcription factor family. MYB33 and MYB65 proteins exhibit 58,4% identity in terms of their amino acid sequence; specifically, they share more than 90% identity in their R2R3 domains and 51% similarity in their carboxy terminal domains (Millar and Gubler 2005). High similarity in aa sequence may indeed suggest some redundancy in function. It was previously shown that MYB33 and MYB65 are involved in anther and pollen development (Dubos et al. 2010; Millar and Gubler 2005). Moreover, it was suggested that both proteins act redundantly in these processes. However, MYB33 and MYB65 are not expressed equally in all plant tissues: MYB33 is expressed in all organs and tissues, with the highest expression of this gene occurring in germinating seeds, mature leaves, flowers, and carpels, while MYB65 is also expressed in all organs; however, the highest amount is detected only in germinating seeds and flowers. Generally, MYB33 is expressed at a higher level than is MYB65, and MYB101 is expressed at the lowest level in all organs, with a slight increase in flowers and seeds during the first stages of germination (Winter et al. 2007). Arabidopsis MYB101, together with MYB97 and MYB120, was shown to function as male factors that control pollen tube–synergid interactions during fertilization (Liang et al. 2013). Together, all these data show that MYBs 33/65/101 have different functions in various processes, possibly by controlling the expression of downstream genes. Moreover, the set of controlled genes may be different depending on the time and place of a given MYB TF action. These data suggest that although all three MYB TFs are expressed in whole plants, the level of their expression varies among the organs studied, which may affect different functions in plant development, strengthening the results obtained in this work. Moreover, these data show the lack of redundancy in their function in plants in response to drought.

The role of MYB TFs from the R2R3 subgroup in the drought response is not confined to the three MYB33/65/101 studied in this work. Previous studies on the *AtMYB44* gene (a member of the R2R3 subgroup) also revealed that this gene plays a role in the plant response to drought (Jung et al. 2008; Wu et al. 2019; Zhao et al. 2018). Plant overexpressing *MYB44* showed enhanced stomatal closure, which provided drought and salinity tolerance. Moreover, it was shown that *AtMYB44* overexpression resulted in reduced expression of genes encoding PP2C phosphatases that are known to be negative regulators of ABA signaling. Maize transgenic plants overexpressing *OsMYB55*, which also belongs to the R2R3 subgroup, also exhibited enhanced tolerance to water deficiency (Casaretto et al. 2016). Overexpression of the potato MYB TF *StMYB1R-1*, which belongs to another subgroup of MYB transcription factors (R1), also resulted in improved plant responses to drought and relatively rapid stomatal closure under drought (Shin et al. 2011). It was shown that StMYB1R-1 enhanced the expression of genes involved in the regulation of water loss. It is highly probable that the mode of action of MYB33/65/101 in the Arabidopsis and potato responses to drought is involved in the same or other pathways that negatively affect ABA signaling and the regulation of water loss.

Arabidopsis MYB33, MYB65 and MYB101 were also shown to inhibit cell division in vegetative plant tissues and thus inhibit plant growth (Allen et al. 2007; Alonso-Peral et al. 2010). The most profound phenotypic effect was found in the case of a mutant carrying an inactivated miRNA159 gene. Our studies on transgenic potato plants harboring overexpressed potato MYB33 and MYB65 revealed a strong dwarf phenotype. Why we did not observe this phenotype in the case of potato and Arabidopsis plants overexpressing Arabidopsis *MYB33, MYB65* or *MYB101* is not clear. The amount of individual overexpressed proteins for each transgenic line may play a crucial role here.

Drought limits the productivity of crop plants. Our results point to the select MYB TFs being interesting, evolutionarily conserved targets for genetic manipulations aiming to obtain crop plants with improved plant tolerance to drought. In potato, tubers are sink organs, which are the most important agronomic potato trait. Damage to the photosynthetic apparatus and changes in photosynthetic processes as a result of drought result in decreasing tuber yield and tuber quality. It will be important to show that the overexpression of the *MYB33, MYB65* and *MYB101* genes is a potential way to increase potato yields under drought. It will also be important to identify target genes acting downstream of these MYB genes. Such an approach will allow the selection of genes with a narrower range of functions than MYB transcription factors have and the avoidance of undesired “off traits” in crop plants.

## Acknowledgments

This work was funded by the National Center of Sciences (NCN); OPUS grant UMO-2012/05/B/NZ9/03383; Preludium grant UMO-2014/15/N/NZ1/00498; Maestro grant 2013/10/A/NZ1/00557; and the KNOW RNA Research Centre in Poznan (01KNOW2/2014). IHAR-PIB Mlochow was funded by Statutory Grant 1-3-00-1-01 from the Polish Ministry of Science and Higher Education.

We would like to thank Prof. Anthony Millar, Australian National University, for enabling us to use and modify genetic constructs containing MYB33 and MYB101 sequences, which were produced in his lab. We would also like to thank Dr. Kaja Milanowska and Marek Zywicki for helping with the bioinformatic analysis, Prof. Magdalena Krzeslowska for consultation on the cuticle analysis and to Zuzanna Jankun for her contribution by genotyping the AthMYB101OE mutants.

## Author contributions

AW carried out the majority of experiments including mutant plant construction and drought experiments; DB took part in the discussion, prepared all figures and calculated statistics; PP took part in the ABA - dependent stomata closure experiments and in the manuscript preparation; DSK participated in evaluation of RWC tests in the case of potato plants, IWF participated in receiving of potato plant material; WM participated in discussions and manuscript preparation, AJ took part in the experiment design and manuscript preparation; ZSK created the concept of the work, designed experiments and wrote the manuscript

---

**Figure S1.**
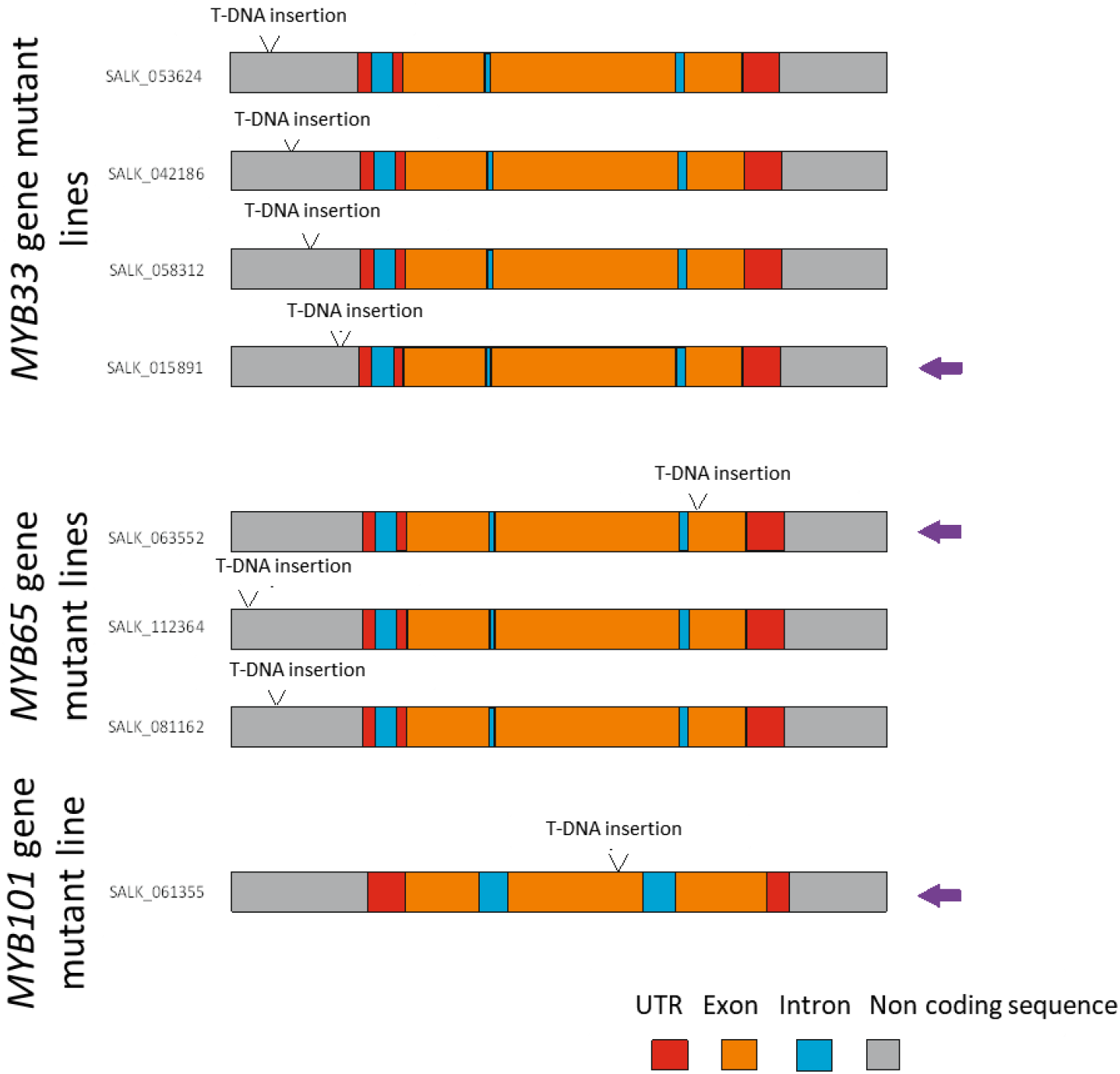
Arabidopsis *MYB33, MYB65* and *MYB101* gene T-DNA insertion lines for analysis of gene knock-down or downregulation. Schematic graphs represent *MYB33, MYB65* and *MYB101*gene structures. Grey color represent gene promotor and 3’ flanking region, red color-5’ and 3’ UTRs, orange color-exons, and blue color-intron. SALK numbers are given at the left side of each gene harboring T-DNA insertion. Localization of T-DNA insertion is marked in the each gene structure. Violet arrows point T-DNA insertion lines for further analysis.

**Figure S2.**
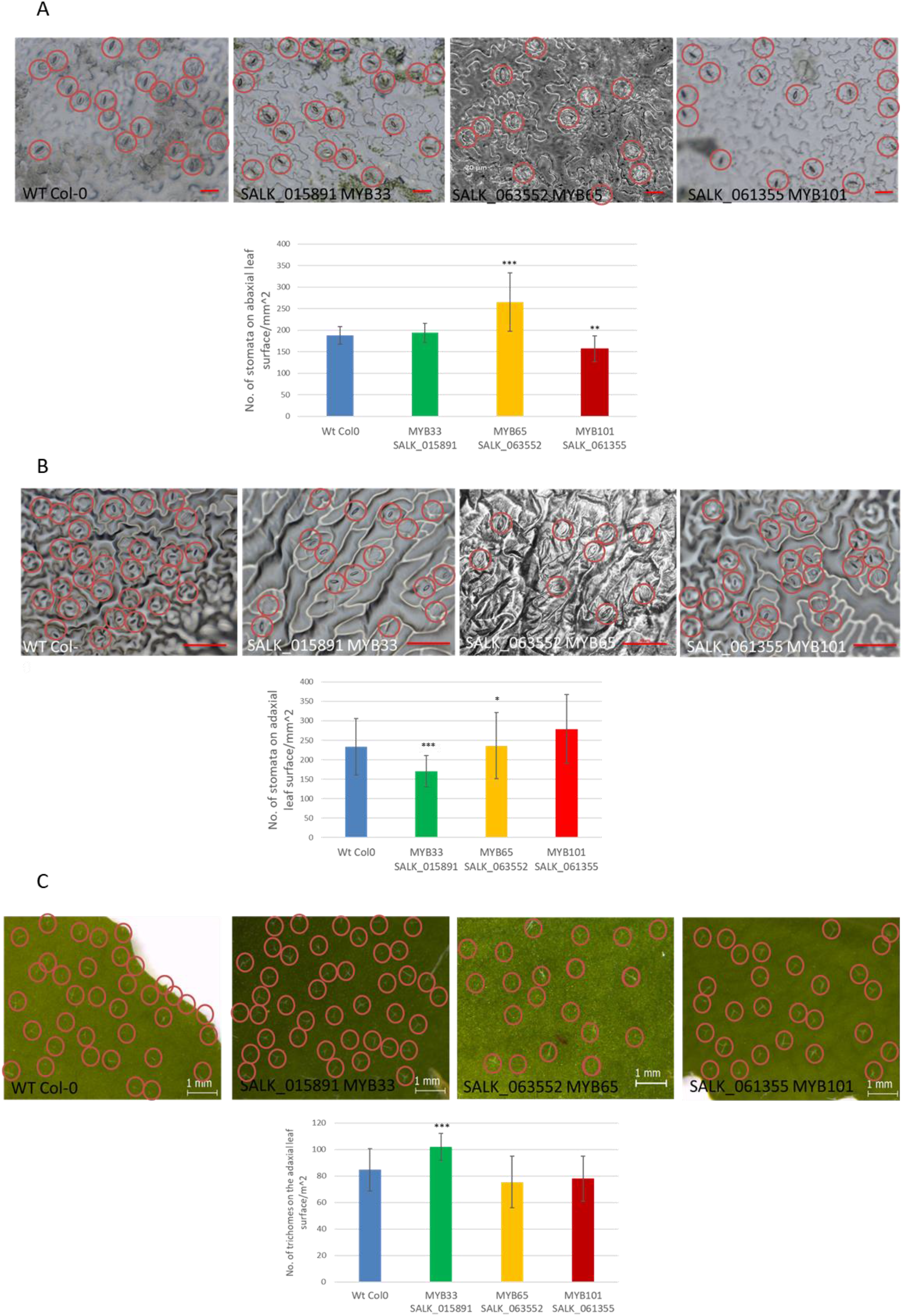
Stomata and trichomes density on the surfaces of Arabidopsis leaves are differentially affected in the Arabidopsis mutant plants with the down regulated *MYB33, MYB65* or *MYB101* expression. Stomata density on both abaxial **(A)** and adaxial **(B)** leaf surfaces in Arabidopsis mutant plants with downregulated level of *MYB33, MYB65* or *MYB101* genes when compared to wild type plants. (A) Upper panel shows light micrographs of abaxial leaf surface. Each stomata is circled in red for visualization. Scale bar – 20 µm. Lower panel presents a graph showing a comparison of the abaxial leaf stomata density in wild type and transgenic plants representing three independent SALK mutant lines exhibiting downregulation of *MYB33, MYB65*, and *MYB101* gene expression, respectively. Blue bar – wild type plants. Colored bars represent selected mutant lines. (**B**) the same experiments as in the (A) panel performed for stomata density on the adaxial leaf surface of wild type and *myb33, myb65*, or *myb101* mutant plants. Upper panel scale bar – 50 µm. **(C)** Upper panel - trichome density on adaxial leaf surfaces in Arabidopsis *myb33, myb65*, and *myb101* mutant plants when compared to wild type plants. Each trichome is circled in red for visualization. Scale bar – 1 mm. Lower panel – a graph showing a comparison of the adaxial leaf trichome density in wild type and transgenic plants representing three independent SALK mutant lines exhibiting downregulation of *MYB33, MYB65*, and *MYB101* gene expression. Blue bar – wild type plants. Colored bars represent selected mutant lines. Values are shown as the mean of ±SD (n=10). Mann-Whitney test, *P* value:* *p*<0.05, ** *p*<0.01;*** *p*<0.001.

**Figure S3.**
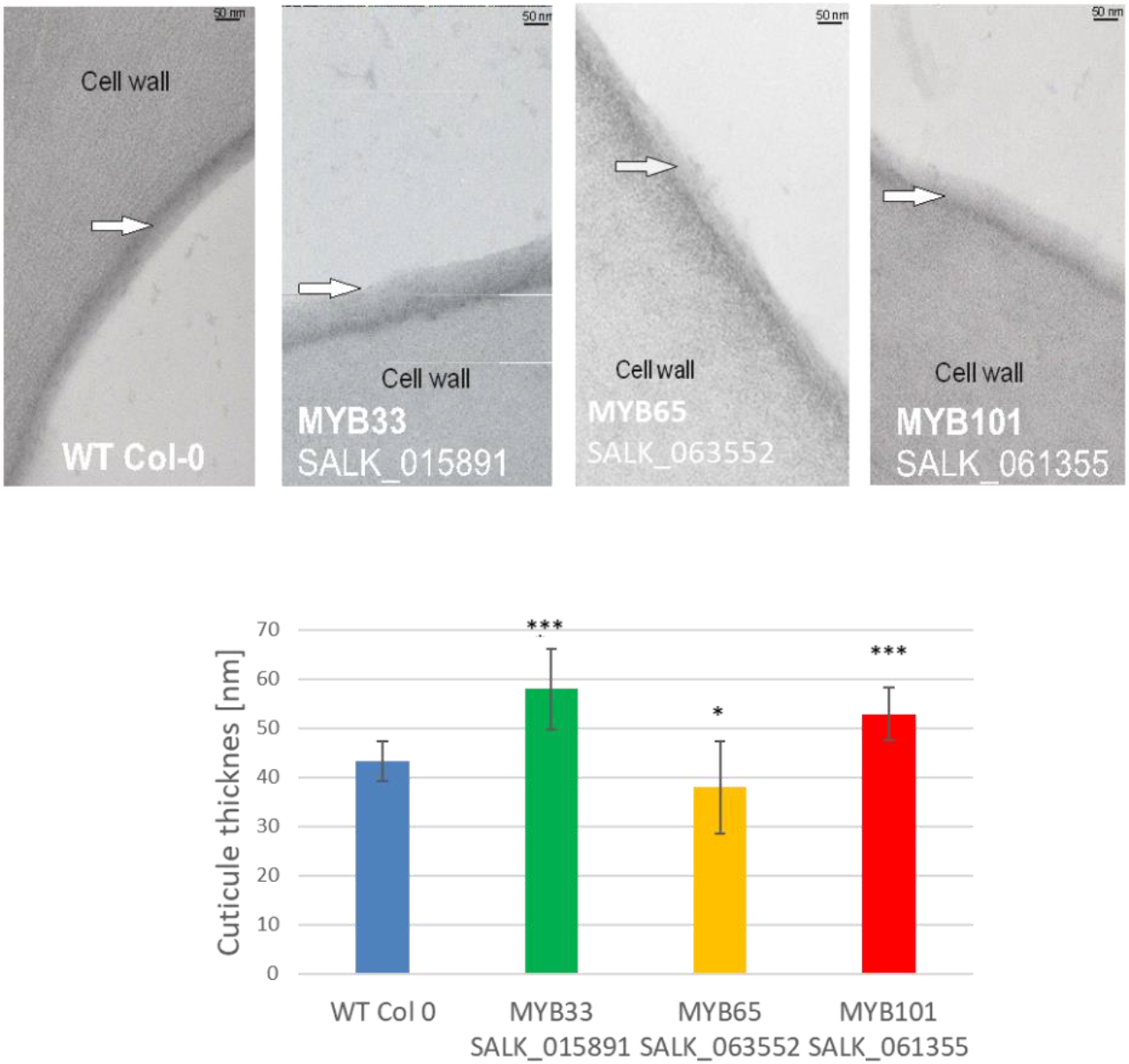
Cuticule ultrastructure is differently affected by the downregulation of *MYB33, MYB65*, and *MYB101* gene expression levels. Adaxial cuticle ultrastructure presented on TEM micrographs (upper panel). Arrows point to the cuticle layer. Scale bar - 50nm. Lower panel - graphs showing measurements of cuticle thickness in each mutant in comparison to WT. Mann-Whitney test, *P* value:* *p*<0,05, ** *p*<0,01;*** *p*<0,001.

**Figure S4.**
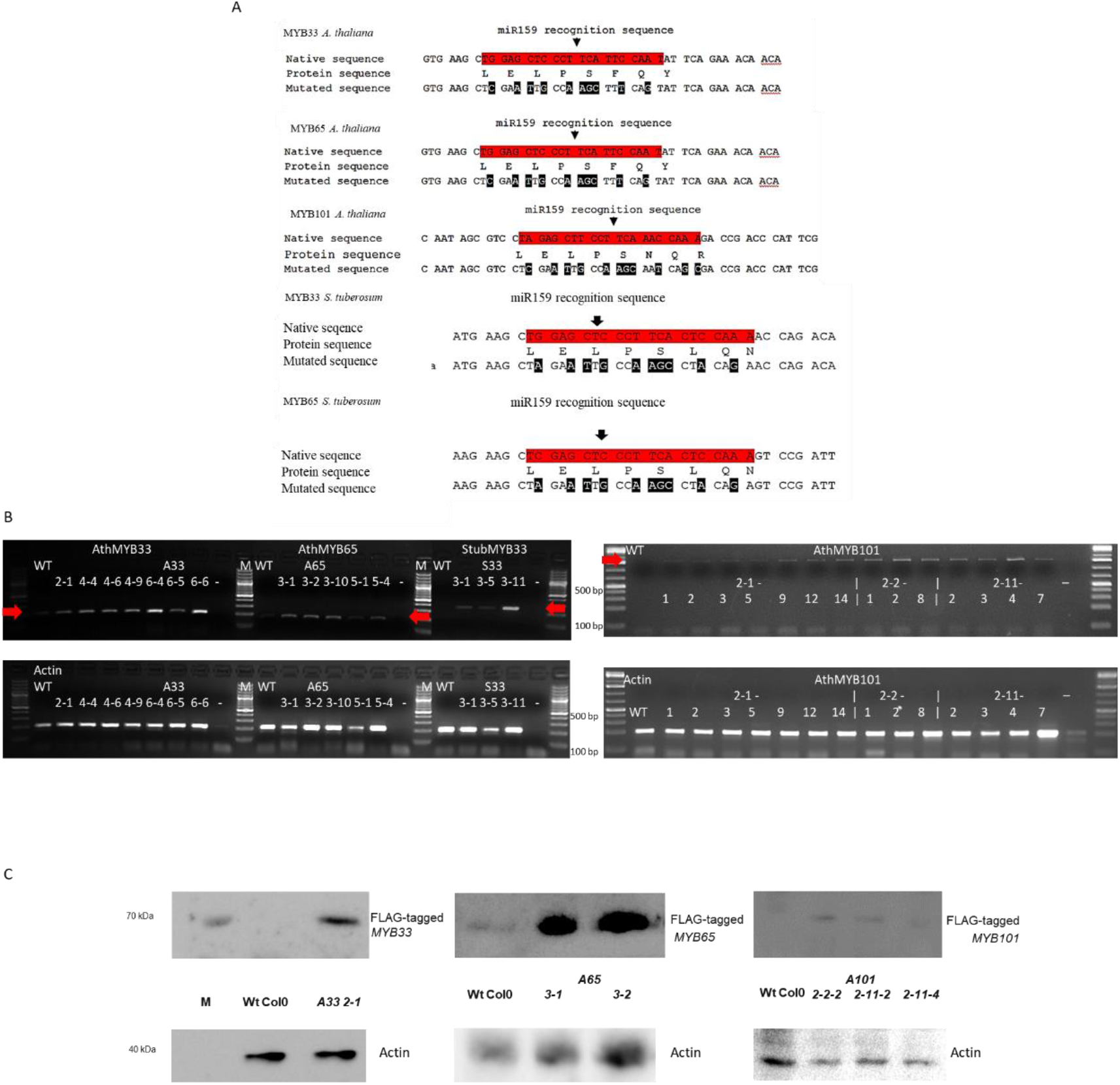
Confirmation of mRNA and protein expression of At*MYB33*, At*MY65*, At*MYB101* and St*MYB33* transgenes in Arabidopsis mutant plants. **A)** Schematic representation of changes in microRNA159 recognition sites in mRNA sequences of studied *MYB* genes. **(B)** Agarose gel electrophoresis of RT-PCR products of *AtMYB33, AtMYB65, AtMYB101*, and *StMYB33* cDNAs in a number of transgenic lines compared to WT plants. Arrows point to the obtained proper products; lower panels show expression of actin cDNA as a control. WT-wild type, M-GeneRuler 100 bp+ DNA Ladder, - negative control. **(C)** Western Blots showing FLAG-tagged MYB proteins expressed in selected Arabidopsis transgenic lines (upper panels) in comparison to WT; anti-actin antibodies were used as a control (lower panels). M – protein weight marker.

**Figure S5.**
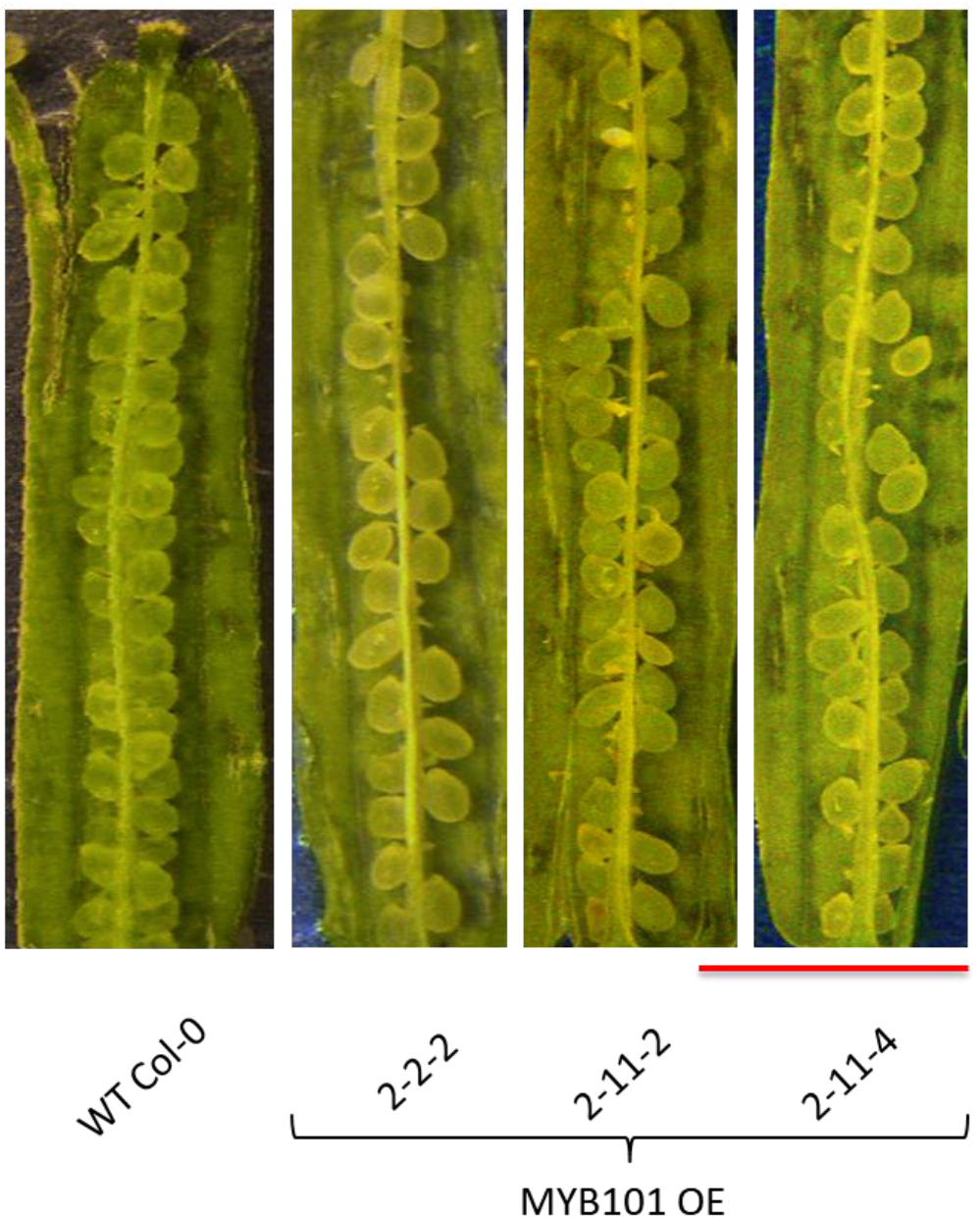
*AtMYB101* OE hemizygote plants siliques reveal embryo lethality of homozygous *AtMYB101*OE seeds. Binocular graphs of Arabidopsis green siliques, cut perpendicularly on one side of the replum, with valves flattened on the both sides of the septum. Three lines: 2-2-2, 2-11-2 and 2-11-4 of plants with overexpression of *AtMYB101*, show spots where seeds did not develop and/or the development was arrested, while in WT plants septum has complete set of the seeds. This result indicates that two copies of *AtMYB101* transgene are lethal for the plants at the embryo developmental stage. Scale bar – 2mm.

**Figure S6.**
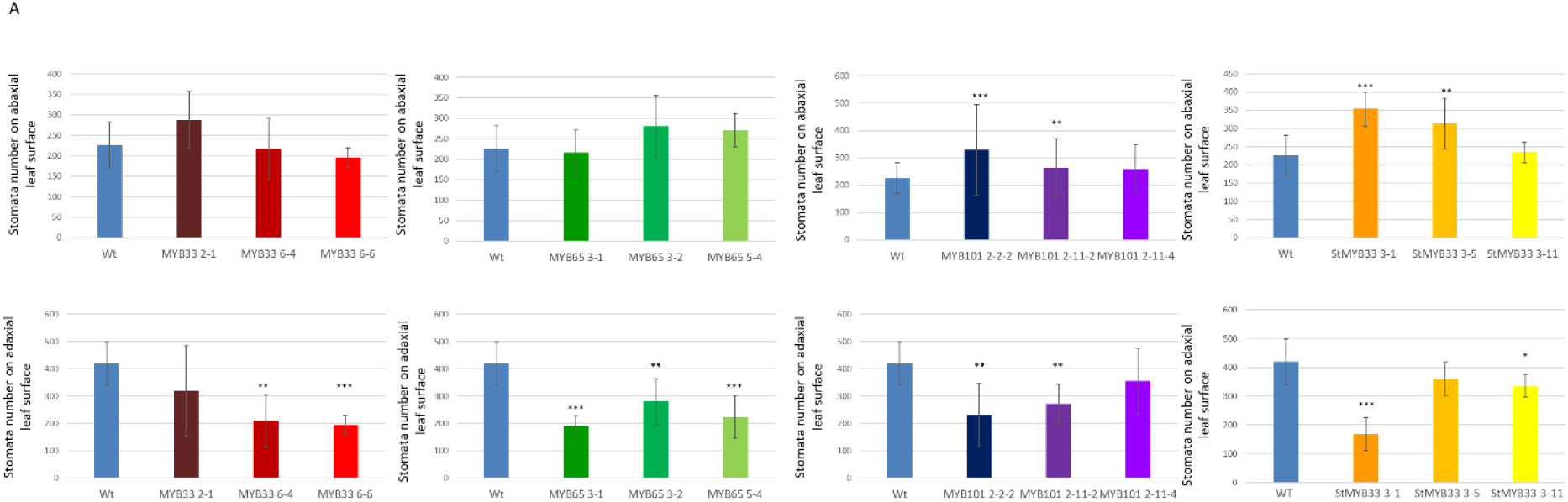
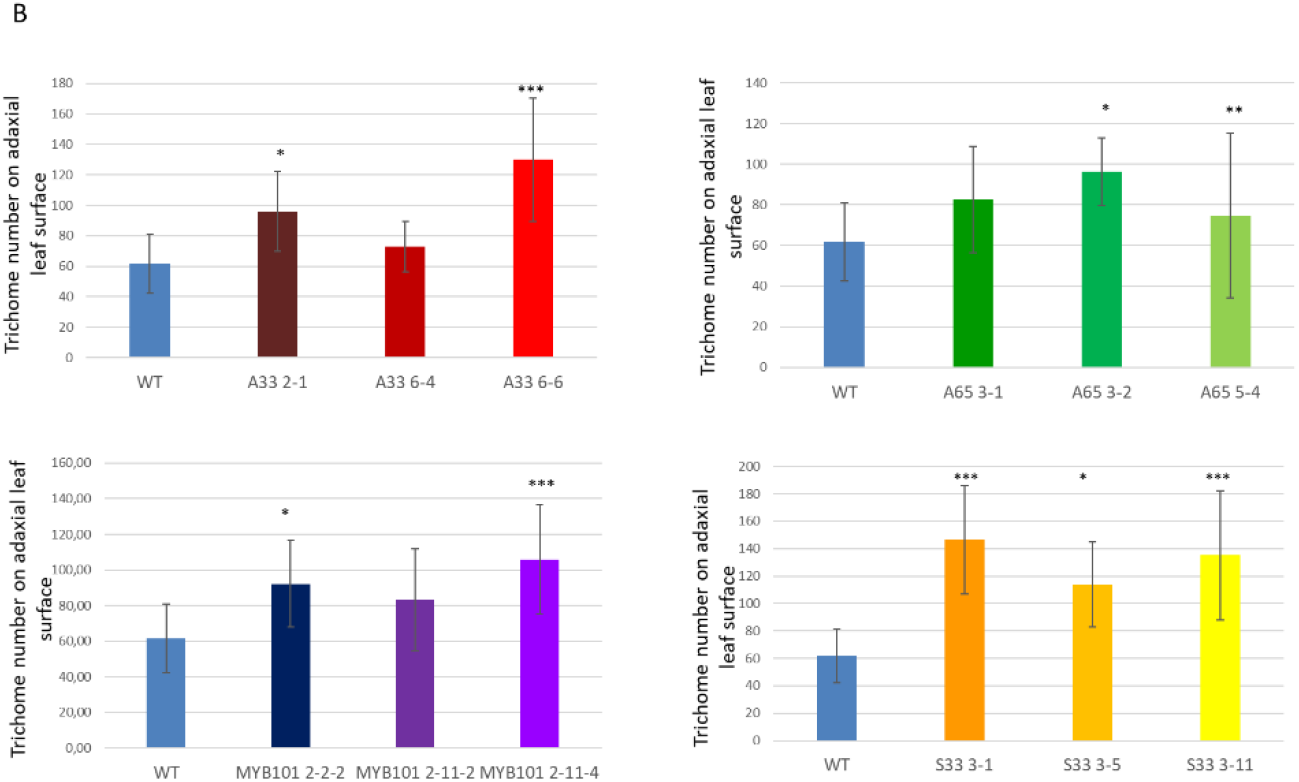
Stomata and trichomes density on the surfaces of Arabidopsis leaves are differentially affected in the Arabidopsis mutant plants overexpressing *AtMYB33, AtMYB65, AtMYB101*, or *StMYB33* transgenes. **A)**Graphs show a comparison of the abaxial (upper panel) or adaxial (lower panel) leaf stomata density in wild type and transgenic plants representing three independent transgenic lines exhibiting overexpression of *MYB33, MYB65*, and *MYB101* gene expression, respectively. Blue bar – wild type plants. Colored bars represent selected mutant lines. (B) Graphs show trichome density on adaxial leaf surfaces in the same Arabidopsis mutant plants as in (A). Values are shown as the mean of ±SD (n=9). Mann-Whitney test, *P* value:* *p*<0.05, ** *p*<0.01;*** *p*<0.001.

**Figure S7.**
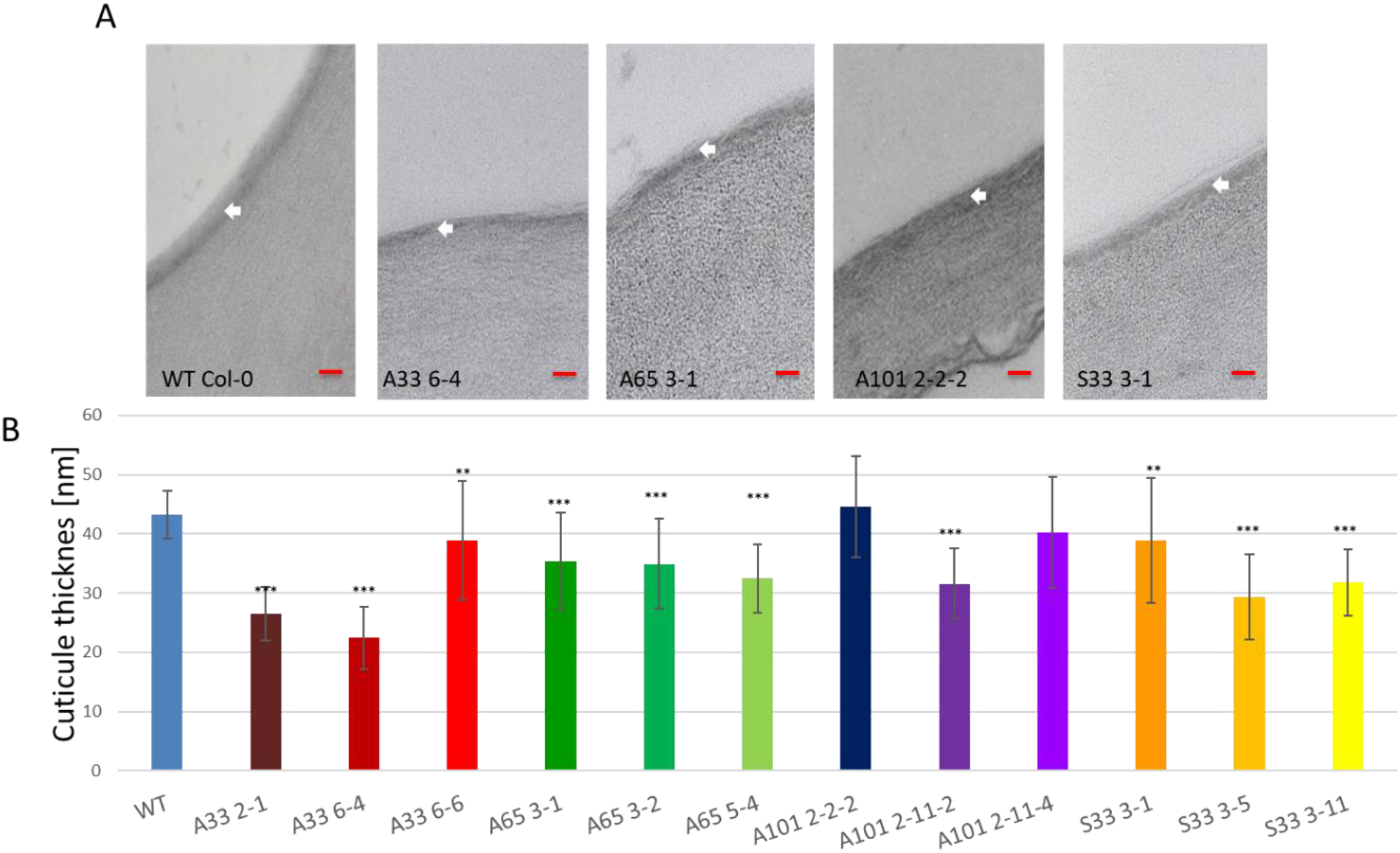
Cuticule is thinner in the transgenic Arabidopsis plants overexpressing *MYB33, MYB65*, and *MYB101* genes. (A) Adaxial cuticle ultrastructure presented on TEM micrographs. Arrows point to the cuticle layer. Scale bar – 50nm. (**B**) Graphs show measurements of cuticule thickness in each mutant in comparison to WT. Mann-Whitney test, *P* value:* *p*<0.05, ** *p*<0.01;*** *p*<0.001.

**Figure S8.**
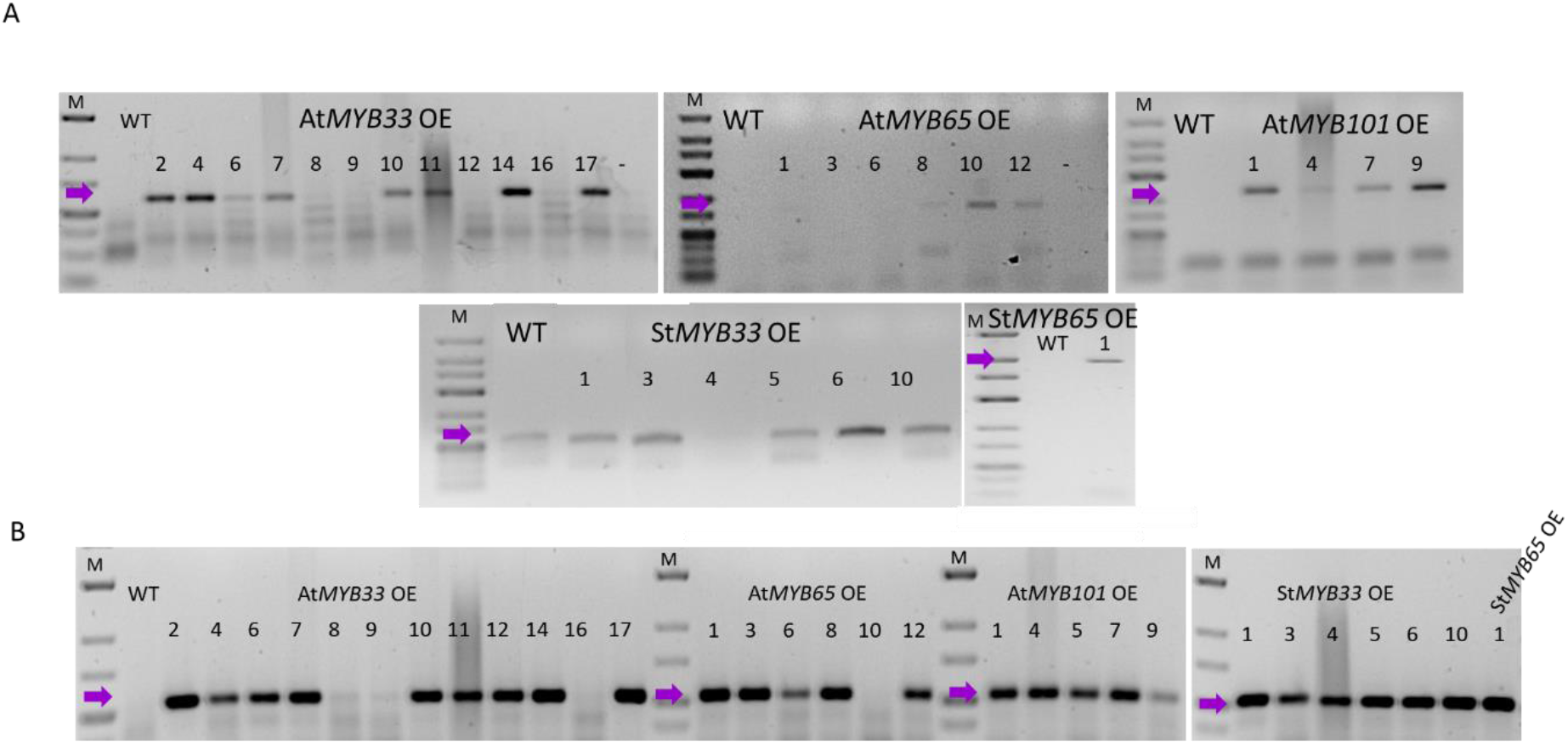
Confirmation of mRNA expression of At*MYB33*, At*MY65*, At*MYB101*, St*MYB33* and *StMYB65* transgenes in potato mutant plants. Agarose gel electrophoresis showing RT-PCR products of fragments of over-expressed cDNAs, respectively (**A**) and hygromycin B phosphotransferase gene mRNA (**B**) in individual transgenic lines of each OE construct, comparing to WT Desiree plants. Numbers represent individual transgenic lines. Arrows point to the obtained proper products. WT-wild type Desiree, M-Low Range GeneRuler DNA Ladder, - negative control.

**Figure S9.**
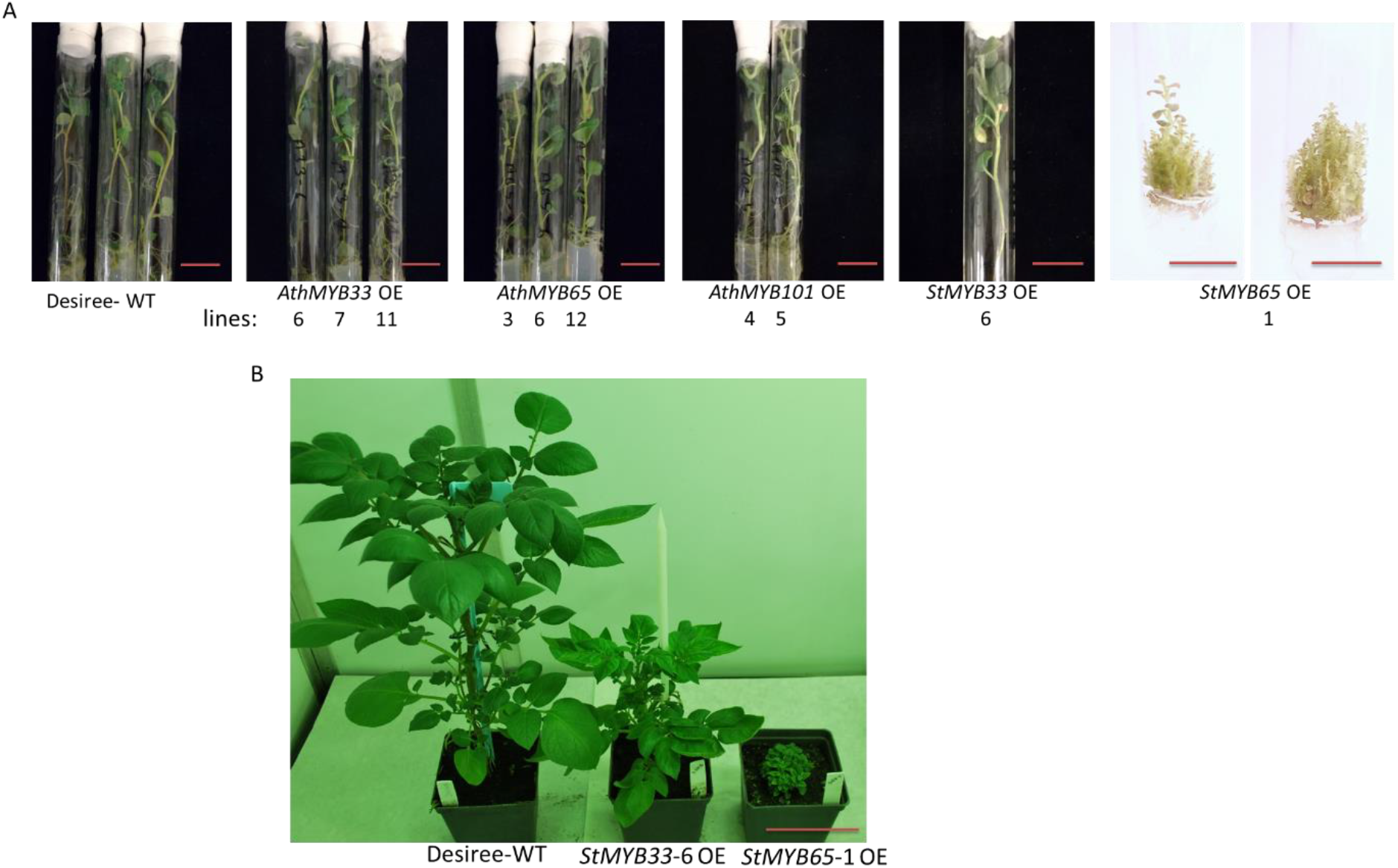
Potato transgenic plants overexpressing *AthMYB33, AthMYB65, AthMYB101, StMYB33* and *StMYB65*, respectively grown in vitro and in pots. (A) In vitro grown potato plants with over expression of *AtMYB33*, AtMYB65, and AtMYB101 genes do not show phenotypic differences in comparison to wild type plants. Only the plants with over expression of *StMYB65* show strong dwarfing phenotype. Scale bars – 2.5cm. (B) Overexpression of potato MYB TFs strongly affects potato phenotype. Comparison of wild type potato plant with *StMYB33* OE and *StMYB65* OE. Scale bar – 15cm.

**Figure S10.**
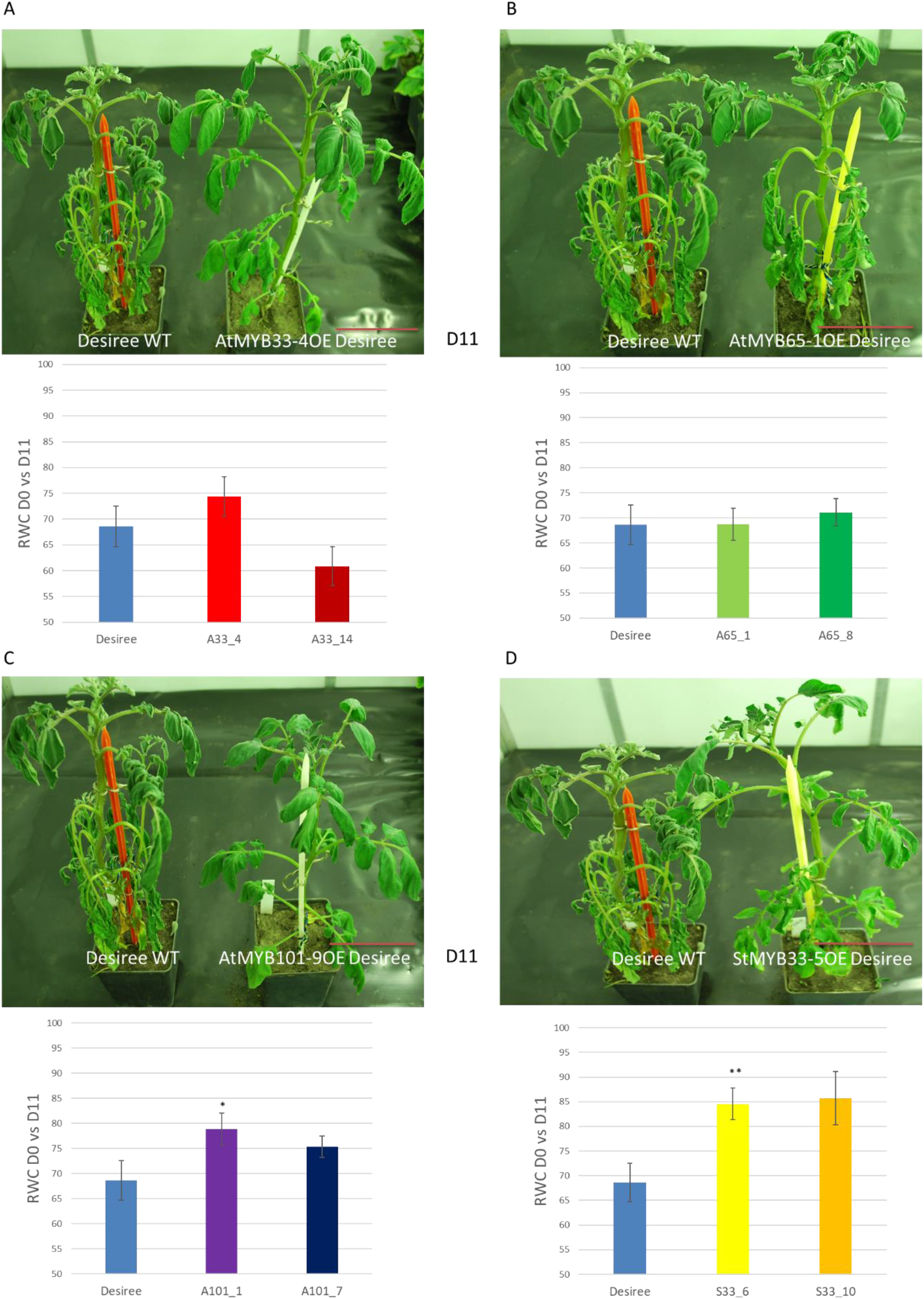
Transgenic lines of potato cultivar Desiree with overexpression of Ath*MYB33*, Ath*MY65*, Ath*MYB101* genes and St*MYB33* gene are resistant to drought. (**A**) Upper panel presents transgenic potato plant overexpressing *AtMYB33* gene (line AtMYB33-4 OE) compared with WT plant. Plants are shown in day 11^th^ (D 11) after water cessation. Lower panel -RWC measurements in leaves from WT and transgenic *AtMYB33* OE (A33) plants after drought stress. The RWC value of the control (Desiree-0,68) is the value of difference between its water content on the D0 and on the D11. Blue bars – WT plants, other colored bars represent potato plants from independent transgenic lines overexpressing *AtMYB33*. (**B, C** and **D**) show the same data obtained for potato transgenic plants overexpressing *AtMYB65* (**B**), *AtMYB101* (**C**), and *StMYB33* (**D**), respectively. Figure descriptions – as in the (A) panels. (A, B, C, and D) RWC data shown as the mean± SD of n=3 independent experiments. Mann-Whitney test, *P* value:* *p*<0.05;** *p*<0.01;*** *p*<0.001. Scale bar – 15 cm

**Figure S11.**
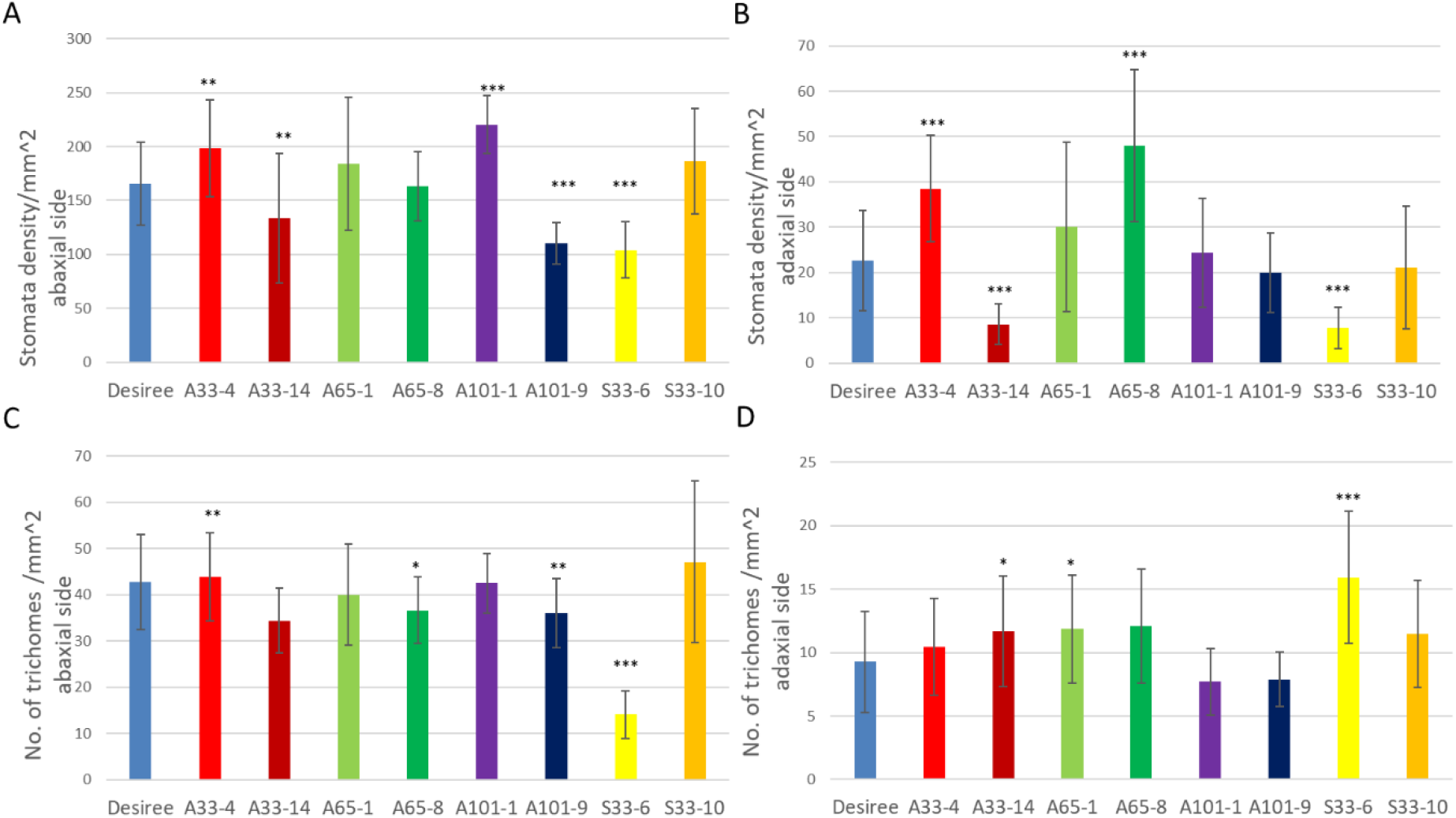
Stomata and trichomes density on the surfaces of leaves are differentially affected in the potato mutant plants overexpressing *AtMYB33, AtMYB65, AtMYB101*, or *StMYB33* transgenes. (A) Graphs show a comparison of the abaxial (A) or adaxial (B) leaf stomata density in wild type and transgenic plants representing three independent transgenic lines exhibiting overexpression of *MYB33, MYB65, MYB101*, and *StMYB33* genes. Blue bar – wild type plants. Colored bars represent selected mutant lines. (C, D) Tables show trichome density on abaxial (C) and adaxial (D) leaf surfaces in the same potato mutant plants as in (A) Values are shown as the mean of ±SD (n=9). *P* value:* *p*<0.05, ** *p*<0.01; *** *p*<0.001; Mann-Whitney test.

